# *In vivo* isogenic modelling unveils *TP53*-associated relapse trajectories in T-cell acute lymphoblastic leukemia

**DOI:** 10.64898/2026.01.19.700255

**Authors:** Stéphanie Gachet, Samuel Quentin, Lucie Hernandez, Loïc Maillard, Marie Passet, Rathana Kim, Hugo Bergugnat, Melha Benlebna, Rémy Jelin, Marine Aglave, Maxime Boy, Véronique Parietti, Pierre Fenaux, André Baruchel, Hervé Dombret, Nicolas Boissel, François Sigaux, Hugues de Thé, Emmanuelle Clappier, Jean Soulier

## Abstract

Many patients with T-cell acute lymphoblastic leukemia (T-ALL) relapse into a treatment-resistant disease. The mechanisms driving relapse remain largely elusive, in part due to the lack of faithful experimental models. Here, we leveraged patient-derived xenograft (PDX) pairs generated from diagnosis and relapse T-ALLs to functionally address the cellular mechanisms driving *TP53-*altered relapse. Beyond inter-T-ALL variability, comparative analyses revealed a unique, cell-intrinsic relapse phenotype that includes greater leukemia-initiating capacity and that can be conferred to diagnosis cells by *TP53* silencing. Transcriptomic profiling linked the relapse phenotype to deregulated OXPHOS metabolism and MYC signaling. Integration of single-cell profilings uncovered *TP53-*wildtype cell populations at diagnosis expressing a relapse profile, possibly reflecting a pre-existing modulation of *TP53* signaling. These cells sequentially evolved towards biallelic *TP53* inactivation at relapse. Collectively, our findings support a model in which T-ALL relapses emerge from a selected pre-existing transcriptional state characterized by deregulated metabolism that favors subsequent *TP53* inactivation.

## INTRODUCTION

T-cell acute lymphoblastic leukemia (T-ALL) is an aggressive hematologic malignancy that represents 15% of pediatric and 25% of all adult ALL cases^1,2^. Multistep deregulation of key transcription factors and genes controlling self-renewal, senescence, apoptosis, differentiation or proliferation in a T-cell precursor leads to clonal transformation and expansion of blast cells in the lymphoid organs and the bone marrow^1,3–7^. Although most T-ALL patients at diagnosis achieve remission with intensive chemotherapy and glucocorticoids, 15% to 40% of the patients subsequently experience relapse into a resistant leukemia with dismal prognosis^8–10^. Therefore, comprehensive understanding of the functional mechanisms underlying relapse and resistance is a major unmet need to develop more efficient therapeutic strategies. While exhaustive genetic and immunophenotypic landscape studies have enabled to identify a large variety of T-ALL subtypes^2,7,11,12^, these last only partially correlate with the onset of relapse. It rather appears that resistance initiates upon treatment by the selection of residual T-ALL cells with cellular features driving cell survival despite therapy and allowing re-expansion^13–19^. Selection of *NT5C2* somatic mutations leading to cell resistance to 6-mercaptopurine is a well-established, although rare, example of resistance-driven mechanism in T- and B-ALL patients^20–24^, but alternative mechanisms remain elusive.

Here, we investigated *TP53* alterations (*TP53*^alt^, i.e. *TP53* somatic mutations and/or deletions) as a factor of resistance acquisition in T-ALL. While rarely detected at diagnosis (2-3%), *TP53*^alt^ are frequently found at relapse (up to 25%) and are associated to poor clinical outcomes^25–30^. *TP53*^alt^ cells are genetically trackable, allowing to detect and characterize them at various stages of T-ALL progression (diagnosis, remission or relapse). Using *in vivo* isogenic patient-derived xenografts (PDX) of diagnosis and relapse T-ALL pairs, we unveiled a unique relapse phenotype that included increased leukemia initiating activity and modulation of key signaling and metabolic pathways such as MYC and OXPHOS. Integration of multi-omics and functional data, including single-cell analyses, allowed us to delineate the relapse trajectory of rare, therapy-selected cells pre-existing at diagnosis that subsequently expanded sequentially and drove relapse.

## RESULTS

### Isogenic, patient-derived xenografts (PDXs) faithfully capture T-ALL diagnosis and *TP53*^alt^-driven relapse genomic patterns

We aimed at uncovering functional mechanisms involved in leukemia resistance and relapse. We focused on *TP53-*altered T-ALL since *TP53* alterations are well-recognized, genetically-trackable, drivers of relapse but are still poorly mechanistically understood in T-ALL^25–29,31^. We selected four cases of T-ALL with *TP53* alterations at relapse only for which cryopreserved diagnosis and relapse patient samples were available. To overcome the limitations of *ex vivo* functional studies on primary samples or cell lines, we established PDXs of paired diagnosis (Dx) and relapse (Rel) leukemia in immunodeficient NOD.Cg-*Prkdc_scid_ IL2rg_tm1Wjl_*/SzJ (NSG) mice (**Figure 1A**). Genomic analyses of the resulting isogenic, Dx-Rel PDX pairs showed that these PDXs largely captured the genomic pattern of the patient cells, both at diagnosis and at relapse, respectively (**Figure 1B** and **Supplementary Tables 1-2**). *TP53* alterations in patient T-ALL relapse cells were missense mutations in the *TP53* DNA-binding domain within mutational hotspot residues R175 and R248 (cases TALL42; and TALL11 and TALL43, respectively) and a less classical but already reported pathogenic mutation H214 (case TALL30) (**Figure 1C**). The other allele was lost by 17p deletion in 3 cases (TALL30, TALL42 and TALL43) or by 17p uniparental disomy of the mutated allele in 1 case (TALL11) (**Figure 1B**). The resulting PDXs carried the original *TP53* missense mutations in association with the loss of the remaining wild-type (WT) allele, as in primary patient samples, or in 1 case only the deleted allele (PDX43R).

**Figure 1.**
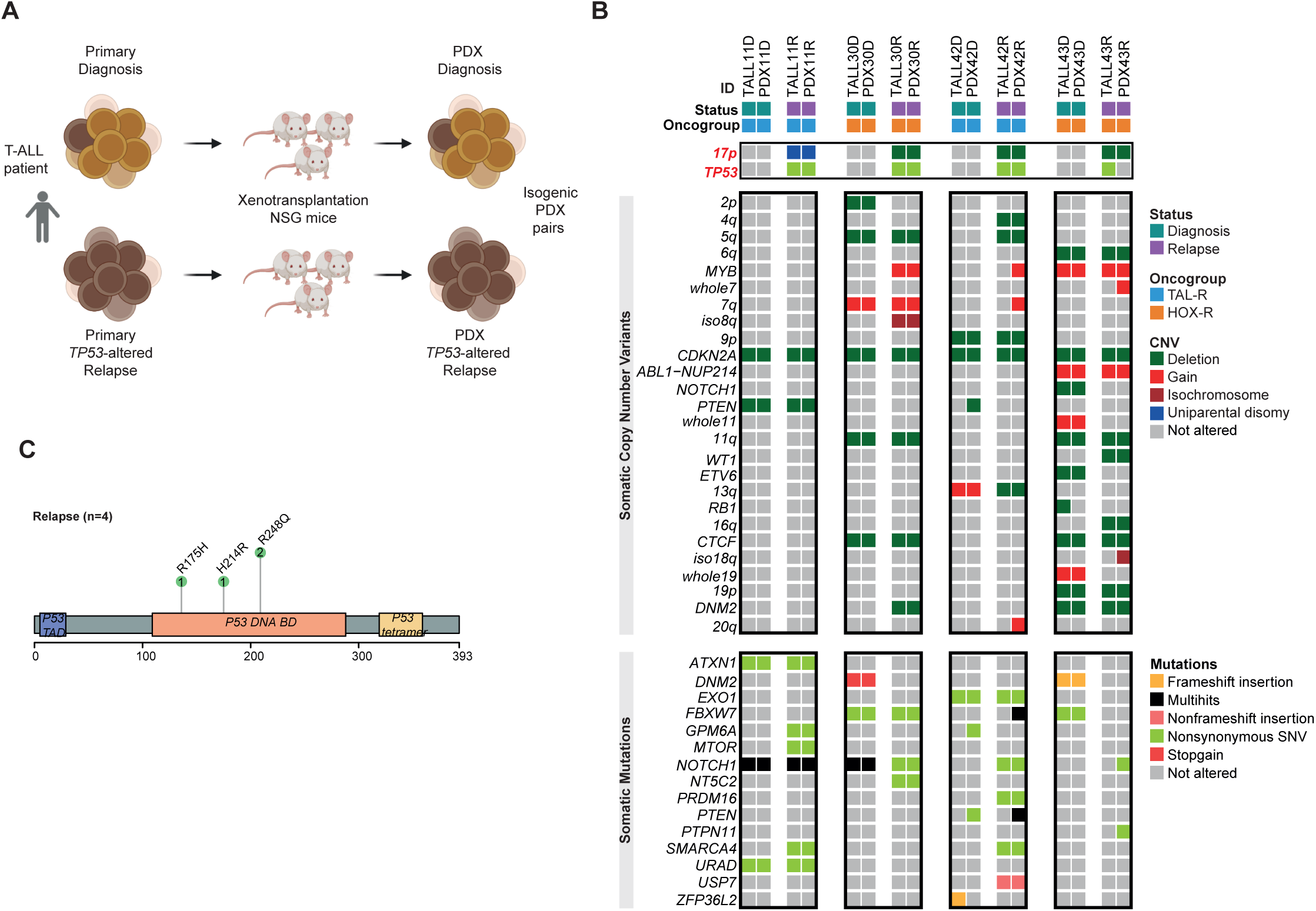
*In vivo* isogenic modelling of diagnosis and relapse *TP53*-altered T-ALL. **A.** Experimental design (Created in BioRender. Gachet, S. (2026) https://BioRender.com/zczr8er). Isogenic Dx-Rel PDX pairs were established after injection into immunodeficient NSG mice of paired primary diagnosis and *TP53*-altered (*TP53*^alt^) relapse samples from the same T-ALL patients. **B.** Comparative Whole Exome Sequencing (WES) analysis of the isogenic Dx-Rel PDX samples established from 4 T-ALL patients who relapsed with *17p/TP53* alterations. In each case, PDX samples (PDXD and PDXR) are compared side-by-side to the corresponding primary samples (TALLD and TALLR, respectively). **C.** Lollipop plot of *TP53* mutations observed in 4 primary relapse samples (TALL11R, TALL30R, TALL42R and TALL43R).

Interestingly, other somatic copy number variants (CNVs) and point mutations (i.e., in addition to *TP53*^alt^) were largely similar in both the diagnosis and relapse samples in the T-ALL pairs, suggesting that *TP53*-driven T-ALL relapse did not acquire stronger genomic instability (**Figure 1B**). Therefore, these T-ALL PDX models faithfully captured the diagnosis and the relapse patient samples, respectively, enabling further functional comparison experiments using this unlimited ressources of isogenic, human leukemic cells.

### Relapse cells display a cell-intrinsic, stable, relapse phenotype compared to their isogenic diagnosis cells

To uncover functional mechanisms involved in leukemia resistance and relapse, we compared the leukemia propagation ability of the diagnosis and relapse T-ALL pairs. First, primary leukemic cells from paired Dx-Rel patient samples were transplanted in parallel at equivalent cell doses in NSG mice, and leukemia onset and mice survival were monitored. Despite strong inter-patient T-ALL heterogeneity, we found in each pair that the relapse cells propagated leukemia more efficiently than their diagnosis counterparts (**Supplementary Figure 1A**), which translated into a significantly shorter mice survival (**Figure 2A** and **Supplementary Figure 1B**). This increased leukemia propagating activity at relapse was observed in all PDXs established from the 4 pairs. We performed multiple iterative transplantations using decreasing cell doses (**Figures 2B-C** and **Supplementary Figures 1C-F**) and found that the propagation phenotype was stable upon serial reinjection, without additional acceleration (**Supplementary Figures 1D ,1F**). Limiting dilution *in vivo* assays (injection of 10^6^ to 1 leukemic cell/mouse) showed that the leukemia initiating cell (LIC) frequencies were increased in the Rel PDXs compared to Dx counterparts (**Figure 2D** and **Supplementary Table 3**). Consistently, *in vitro* long-term culture-initiating cell assays in coculture with MS5-DL1 stromal cells^32^ showed a significantly increased clonogenicity in Rel PDXs from all T-ALL pairs, demonstrating that increased LIC activity is a cell-intrinsic feature of relapse cells (**Supplementary Figure 1G** and **Supplementary Table 3**). Real-time and end-point *in vitro* proliferation using Bromodeoxyuridine (BrdU) pulse labelling and apoptosis experiments, in particular cleaved caspase 3 staining, showed increased proliferation and decreased apoptosis in Rel versus Dx PDX cells (**Figures 2E-F** and **Supplementary Figures 1H-I**). Moreover, homing of the leukemic cells in the bone marrow, spleen and liver at 24 hrs was increased in PDXs from Rel for the 4 T-ALL pairs (**Supplementary Figures 1J-K**). However, when homing was bypassed by direct injection of leukemic cells into the femoral bone marrow cavity, leukemic cells from Rel PDXs still propagated leukemia more efficiently than Dx PDXs (**Supplementary Figures 1L-M**).

**Figure 2.**
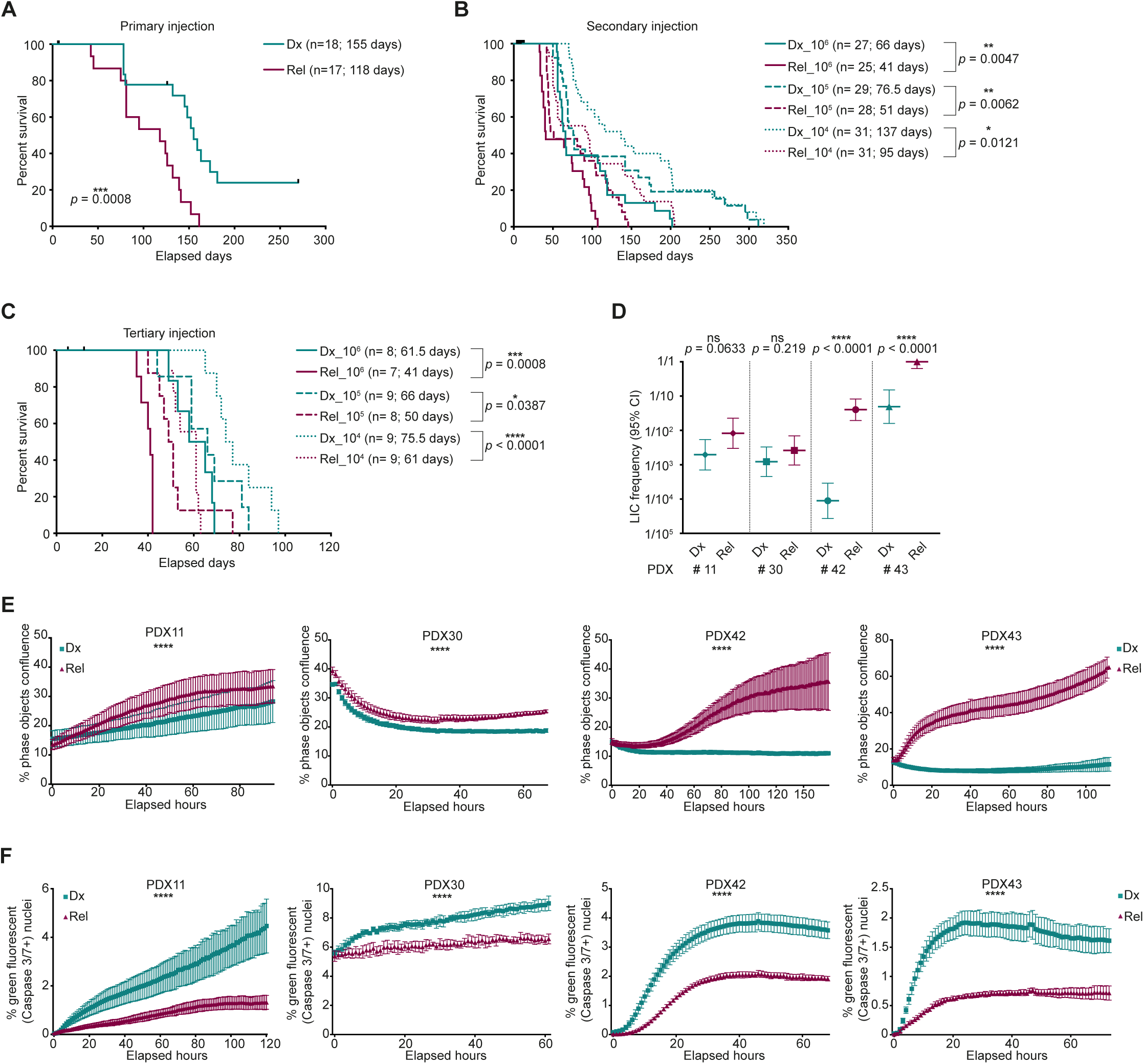
*TP53*^alt^ T-ALL relapse cells display a cell-intrinsic gain of malignancy phenotype. **A.** Kaplan-Meier survival curves of PDX mice engrafted with primary isogenic Dx-Rel leukemic cells from 4 T-ALL patients (TALL11, TALL30, TALL42 and TALL43). N=4-5 recipient mice injected at equivalent cell doses per sample. **B-C.** Parallel serial transplantations of paired Dx-Rel T-ALL PDX samples. Kaplan-Meier survival curves of mice injected with 3 cell doses (10^6^, 10^5^ and 10^4^ cells) of isogenic Dx-Rel PDX pairs (n = 4 TALL cases) raised from primary (**B**) and secondary (**C**) xenografts. **D.** LIC frequencies of isogenic Dx-Rel PDX sample pairs calculated from *in vivo* limiting dilutions. **E-F.** Real-time *in vitro* cell proliferation and apoptosis of leukemic cells from isogenic Dx-Rel PDX pairs, measured with the Incucyte® Live-Cell Analysis System. Proliferation was quantified by measuring the proportion of phase objects confluence over time, without labels **(E)**, and apoptosis by the quantification of green fluorescently labelled nuclei after activated caspase-3/7 cleavage of inert Incucyte® Caspase-3/7 dyes **(F)**. Statistical data in this figure are represented as mean ± s.d. Significance is indicated by *P* values; A-C: Log-rank test (Mantel-Cox); D: overall tests for differences using ELDA software; E-F: two-way Anova test. Number of recipient mice and median survival are indicated into brackets on Kaplan-Meier survival curves. ns, not significant; *, *p* < 0.05; **, *p* < 0.01; ***, *p* < 0.001; ****, *p* < 0.0001.

Collectively, our data indicate that despite inter-T-ALL heterogeneity, *TP53*^alt^ mediate a unique, cell-intrinsic, relapse phenotype that is associated with increased proliferation, cell survival, and cell homing and post-homing advantage, resulting in higher LIC activity.

### Experimental *TP53* silencing confers the cell-intrinsic relapse phenotype to diagnosis leukemic cells

To model *TP53* alterations as driver of the relapse phenotype, we inactivated *TP53* using a lentiviral shRNA approach in Dx PDX leukemic cells followed by transplantation in NSG mice (**Figure 3A** and **Supplementary Figure 2A**). *TP53* downregulation increased the leukemia propagation capacities of Dx PDX cells when transplanted under parallel and competitive settings (**Figure 3B** and **Supplementary Figures 2B, 2D-E**), and resulted in enhanced leukemia spreading and shorter survival of the recipient mice (**Figure 3C** and **Supplementary Figure 2C**). This phenotype was retained in secondary transplants, both in parallel and competitive settings (**Figures 3D-F** and **Supplementary Figures 2F-H**).

**Figure 3.**
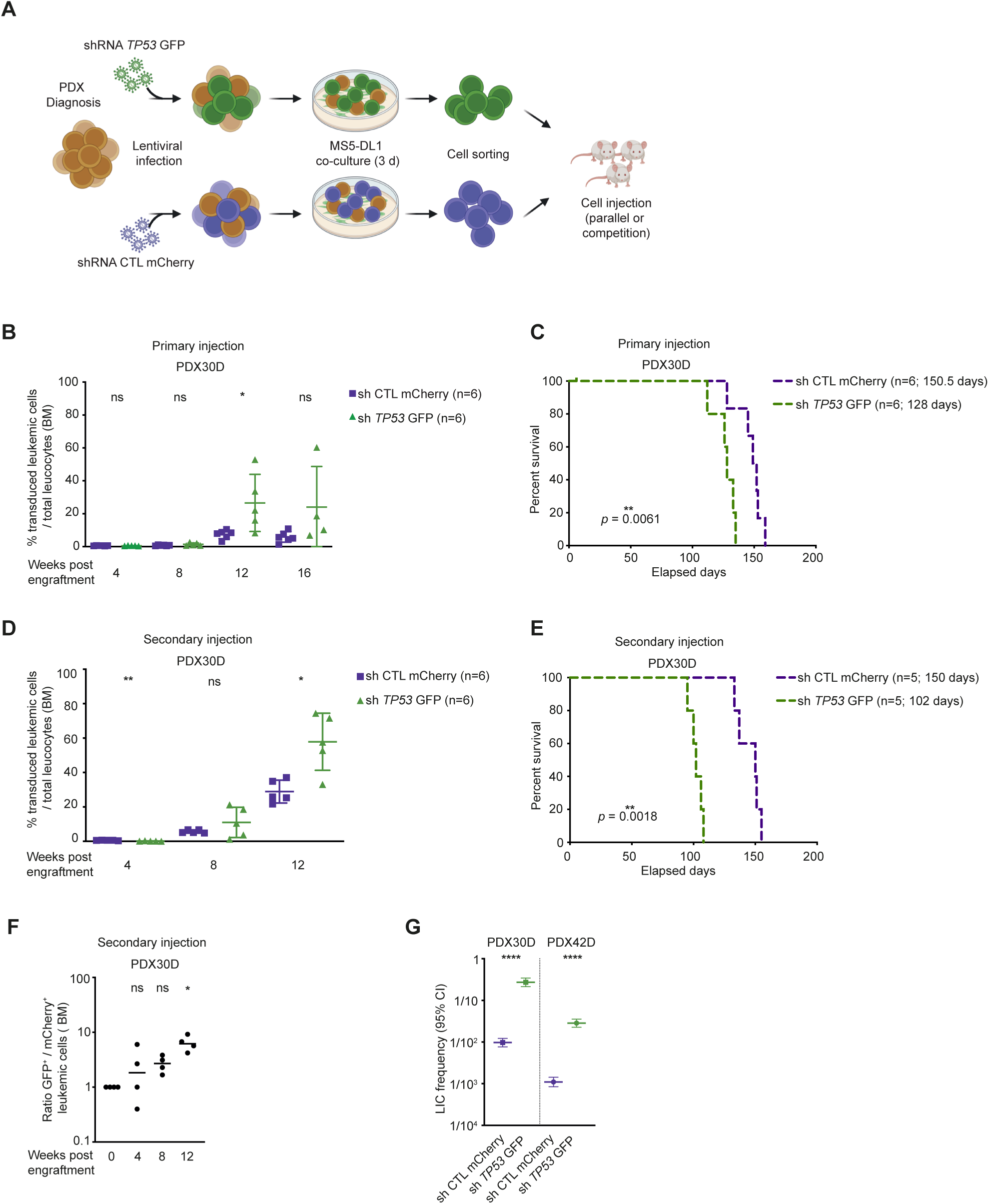
*TP53* silencing confers a relapse phenotype to diagnosis leukemic cells. **A.** Experimental design of *TP53* silencing in diagnosis cells from PDX samples using a lentiviral shRNA approach (Created in BioRender. Gachet, S. (2026) https://BioRender.com/usb6t9c). **B-C.** Primary transplantation of paired control mCherry^+^ (shCTL mCherry) and *TP53* KD GFP^+^ (sh *TP53* GFP) leukemic cells from PDX30D injected in parallel settings into recipient mice. **B.** Engraftment kinetics in the bone marrow (BM) of recipient mice (n=6) injected with paired control and *TP53* KD leukemic cells from PDX30D. **C.** Kaplan-Meier survival curves of mice injected with paired control and *TP53* KD leukemic cells from PDX30D. **D-F.** Secondary transplant of paired control mCherry^+^ (shCTL mCherry) and *TP53* KD GFP^+^ (sh *TP53* GFP) leukemic cells from PDX30D injected either in parallel (panels D-E) or competitive (panel F) settings into recipient mice. **D.** Engraftment kinetics in the BM of recipient mice (n=5) injected in parallel settings with cells from control and *TP53* KD conditions. **E.** Kaplan-Meier survival curves of mice injected in parallel settings with sorted control or *TP53* KD leukemic cells from PDX30D. **F.** Engraftment kinetics in the BM of recipient mice (n=4) injected with sorted cells from both control and *TP53* KD in competitive settings (ratio 1:1). Data are represented as ratios of *TP53* KD GFP^+^ over control mCherry^+^ leukemic cells within BM of the same recipient mouse and compared to the input. **G.** LIC frequencies of paired control-*TP53* KD samples calculated from *in vitro* long-term culture-initiating cell assays of sorted GFP^+^ or mCherry^+^ CD45^+^CD7^+^ leukemic cell from diagnosis PDX samples (PDX30D and PDX42D). LIC frequencies are estimated based on the proportions of positive wells for GFP^+^/mCherry^+^ CD45^+^CD7^+^ leukemic cells evaluated after 2 and 4 weeks of culture. Statistical data in this figure are represented as mean ± s.d. Significance is indicated by *P* values; B and D: Mann-Whitney test; C and E: Log-rank test (Mantel-Cox); F: Kruskal-Wallis test; G: overall tests for differences using ELDA software. Number of recipient mice and median survival are indicated into brackets on Kaplan-Meier survival curves. ns, not significant; *, *p* < 0.05; **, *p* < 0.01; ****, *p* < 0.0001.

*In vitro* long term culture-initiating cell assays and proliferation experiments on the *TP53*-silenced Dx PDX cells confirmed the relapse phenotype, indicating a *TP53*-dependent increased clonogenic activity (**Figure 3G**, **Supplementary Table 3** and **Supplementary Figure 2I**). Finally, *in vivo TP53*-silenced Dx cells also displayed increased homing and post homing capacities (**Supplementary Figures 2J-M**). Altogether, these results demonstrate that *TP53* silencing suffices to confer functional features of the relapse phenotype to patient diagnosis cells.

### *TP53* alterations deregulate MYC signaling and metabolism pathways in T-ALL cells and directly contribute to the *relapse phenotype*

To dig into the biological pathways deregulated by *TP53*^alt^ in T-ALL relapse, we compared transcriptional profiles of isogenic Dx and Rel T-ALL PDX samples. While 91 genes were upregulated at relapse, 181 genes were downregulated including *TP53* itself (except in PDX11R where *TP53* was point-mutated on both alleles) (**Figure 4A** and **Supplementary Figures 3A-B**). Geneset enrichment analysis (GSEA) identified the MYC, E2F, and G2M checkpoint pathways as enriched in Rel cells along with several cell metabolism pathways, including oxidative phosphorylation (OXPHOS), glycolysis and cholesterol homeostasis, defining a relapse expression signature (**Figure 4B**). Moreover, we interrogated genesets linked to cell proliferation, invasiveness and stemness, and evidenced their upregulation at relapse (**Supplementary Figure 3C**). Analysis in secondary transplant PDXs showed consistent expression patterns, demonstrating functional stability (**Supplementary Figures 3D-G**). The relapse expression signature was also retrieved from the corresponding primary T-ALL samples, further outlining that our Dx-Rel PDX models faithfully captured the features of the patient cells (**Figure 4C**).

**Figure 4.**
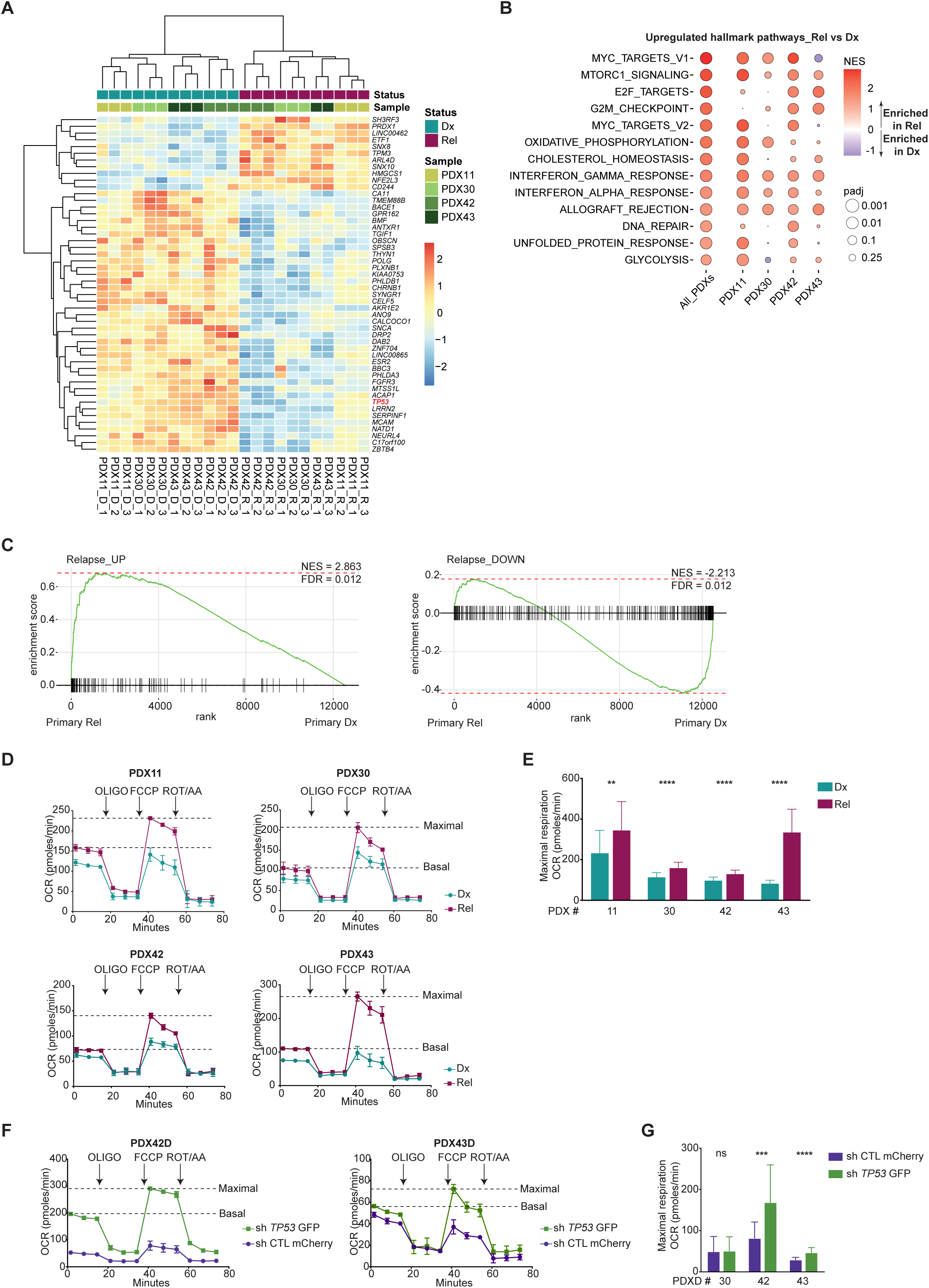
*TP53* alterations drive a relapse phenotype through MYC signaling and metabolism deregulation. **A.** Supervised hierarchical clustering of the top fifty most differentially expressed genes between Dx and Rel T-ALL samples. Heatmap representation of the most variables genes at Dx and Rel in 4 isogenic PDX pairs (primografts). *TP53* expression is highlighted in red. Independent triplicates are analysed for each isogenic PDX pair. **B-C.** GSEA of enriched pathways in *TP53*^alt^ T-ALL relapse samples. **B.** Comparative Gene Set Enrichment Analysis (GSEA) of differentially expressed genes in Rel and Dx samples from 4 isogenic PDX pairs. Representation of selected Hallmark pathways significantly upregulated at relapse, defining a relapse expression signature in all isogenic Dx-Rel PDX pairs pooled together (1^st^ column “All_PDXs”) and in each individual PDX pair (2^nd^ to 5^th^ columns; respectively PDX11, PDX30, PDX42 and PDX43) (*P* adj <0.05). **C.** GSEA of the upregulated (upper panel) and downregulated (lower panel) relapse expression signature obtained from PDX samples (panel B) tested in the original T-ALL patients samples. UP_PDX, upregulated transcriptomic signature in Rel PDXs; DOWN_PDX, downregulated transcriptomic signature in Rel PDXs. FDR, false discovery rate; NES, normalized enrichment score. **D-G.** Direct evaluation of mitochondrial respiration in leukemic cells from isogenic Dx-Rel PDXs using Seahorse assays. **D.** Real-time measurements of oxygen consumption rate (OCR) in isogenic Dx-Rel leukemic cells from 4 T-ALL cases. **E.** Quantification of maximal respiration in leukemic cells from 4 isogenic Dx-Rel PDX pairs evaluated in 3 independent experiments. **F.** Real-time measurements of OCR in *TP53*-silenced (*TP53* KD) and matched control leukemic cells from diagnosis PDX samples (PDX42D and PDX43D). **G.** Quantification of maximal respiration in *TP53* KD and control leukemic cells from 3 independent diagnosis PDX samples (PDX30D, PDX42D and PDX43D) evaluated in 2 independent experiments Statistical data in this figure are represented as mean ± s.d. Significance is indicated by *P* values; E and G: Mann-Whitney test. ns, not significant; **, *p* < 0.01; ***, *p* < 0.001; ****, *p* < 0.0001.

Considering the role of TP53 in regulating MYC signaling and metabolism pathways^33,34^, we focused on OXPHOS and glycolysis and performed real-time measurements of oxygen consumption rate (OCR) and extra cellular acidification rate (ECAR) in isogenic Dx-Rel PDX pairs. We found increased basal and maximal mitochondrial respiration, along with increased glycolysis, in the Rel vs Dx T-ALL cells (**Figures 4D-E** and **Supplementary Figures 3H-K**). Consistently, experimental silencing of *TP53* in the diagnosis PDX cells conferred a relapse mitochondrial and glycolytic profile, demonstrating a driver role of *TP53* inactivation into deregulation of metabolic functions (**Figures 4F-G** and **Supplementary Figures 3L-N**).

### Backtracking pre-relapse *TP53*^alt^ clones at diagnosis

As T-ALL relapse may emerge from minor pre-existing subclones at diagnosis^13,17,19,22,35,36^, we next aimed to backtrack rare *TP53*-mutated cells that would constitute a genetically defined pre-relapse reservoir. We performed ultra-sensitive, quantitative, droplet digital PCR (ddPCR) approach on the isogenic Dx-Rel primary and PDX samples. We retrieved the expected *TP53*-associated relapse mutation in the primary and/or PDX sample from diagnosis in 3 out of 4 cases (all but TALL11), with very low variant allele frequencies (VAF) ranging from 0.01 to 0.04% that could be extrapolated to approximately 1/1250 to 1/5000 total cells, i.e., below the usual medical diagnosis detection thresholds of dominant genetic subclones). These results indicate the presence at diagnosis of a very minor *TP53*-mutant clone that was further selected and expanded in the post-treatment relapse sample. Interestingly, a cryopreserved remission sample was also available in case TALL11, in which we detected the expected *TP53*-R248Q mutation with a low frequency (VAF, 0.02%; approximately 1/2500 total cells), demonstrating the existence of the mutant cells reservoir during treatment. When this remission sample was injected into NSG mice, a full-blown leukemia developed from these rare cells. All the resulting leukemic cells carried the *TP53*-R248Q missense mutation. Interestingly, this leukemia retained a wildtype allele (*TP53*-R248Q allele frequency, 46% in sorted leukemic cells), contrasting with the relapse cells that demonstrated copy-neutral loss of the wildtype allele (*TP53*-R248Q allele frequency, 92%). This highlights the selective advantage upon treatment conferred by the monoallelic point mutation to a minor clone at remission, which further expanded in the PDX. The relapse cells displayed a complete loss of the wildtype *TP53* allele and function with duplication of the *TP53*-R248Q allele through acquisition of an uniparental disomy.

Those data demonstrate that relapse emerged from a pre-existing ultra-minor *TP53*-mutant clone that expanded during treatment through selective growth advantage, and subsequent acquisition of bi-allelic *TP53* loss at full relapse.

### Combined single-cell profiling uncovers pre-relapse, reservoir cells at diagnosis that express features of the relapse phenotype

To further evaluate whether the rare *TP53*-mutant population from diagnosis expressed a relapse-like transcriptional state, we performed single-cell RNAseq analysis on the 4 T-ALL pairs. The 16 resulting samples (8 primary and 8 from PDXs) did not globally cluster by their diagnosis or relapse nature (i.e., all diagnosis *versus* all relapse samples), nor by their origin (i.e., all primary *versus* all PDX samples), or by cell cycle repartition, but rather by individual T-ALL case (**Figure 5A** and **Supplementary Figures 4A-B**), underlying prominent inter T-ALL biological heterogeneity. GSEA confirmed oxidative phosphorylation signature as the most differentially expressed pathway in all relapse samples along with cell proliferation, cell signaling and cell metabolism pathways (**Supplementary Figures 4C-F** and **Figures 5B-C**).

**Figure 5.**
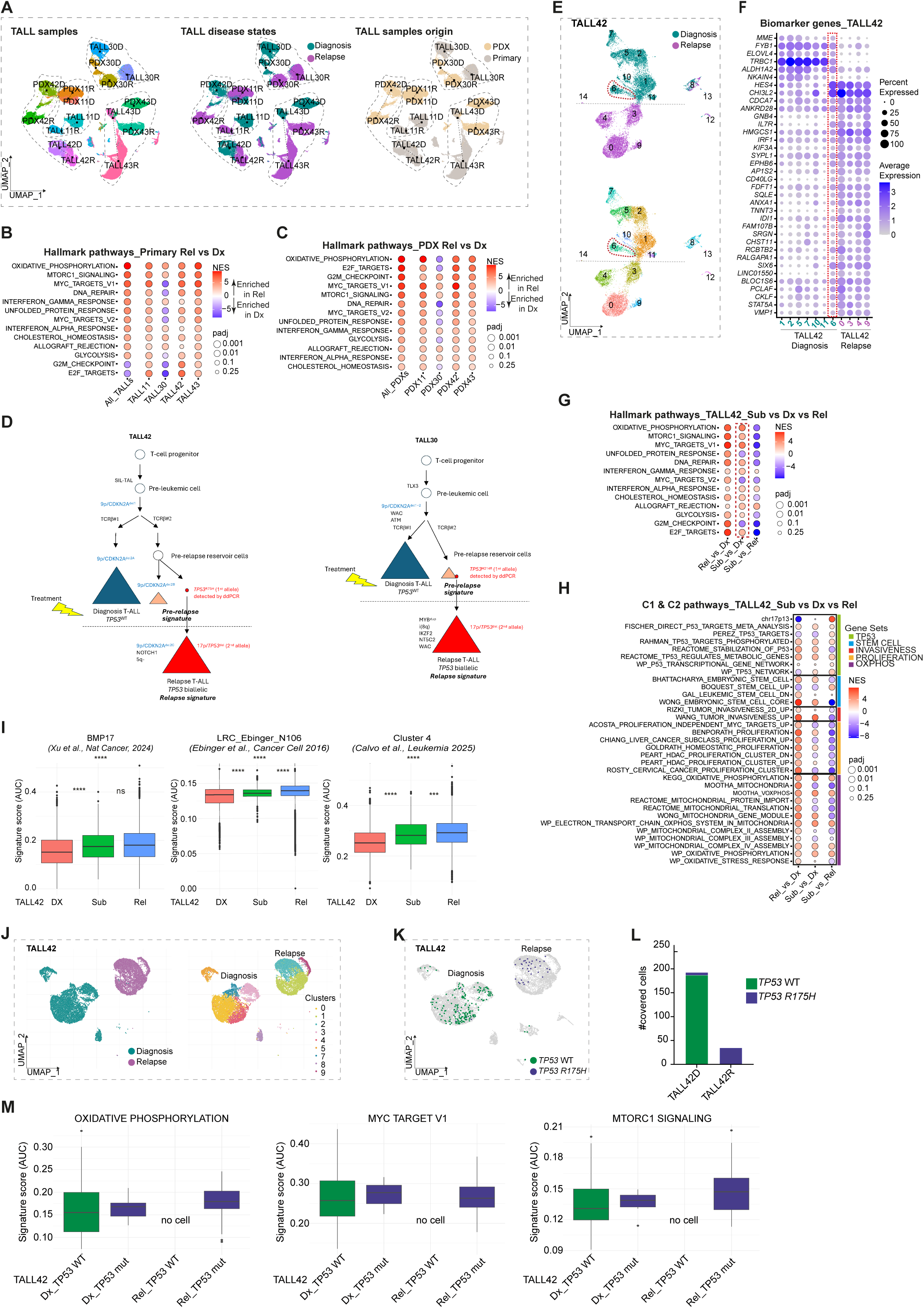
Single-cell profiling of isogenix Dx-Rel T-ALLs can unveil a pre-relapse cells reservoir at diagnosis. **A.** UMAP visualizations of isogenic Dx-Rel primary samples from 4 T-ALL patients and their derived PDXs analysed by short read single-cell RNA (scRNA) sequencing. UMAP plots of 117.119 single cells coloured by T-ALL samples (n=16, primary and PDXs samples from 4 T-ALL cases at diagnosis and relapse; left panel), leukemic cell identity (Diagnosis versus Relapse; middle panel) and by leukemic cell origin (Primary versus PDX; right panel). **B-C.** GSEA of biological pathways enriched at relapse in primary and PDX T-ALL samples. GSEA from pseudo bulk analysis of differentially expressed genes between relapse (Rel) and diagnosis (Dx) samples from isogenic patient (**B**) and PDX (**C**) T-ALL pairs retrieves the relapse expression signature defined in Figure 4B. Representation of selected Hallmark pathways significantly deregulated at relapse (*P* adj <0.05) in all isogenic pairs pooled together (1^st^ column “) and in each individual pair (2^nd^ to 5^th^ columns). Note that TALL30 case eventually diverged from the other cases by the late acquisition of a *NT5C2* mutation, known to drive resistance to maintenance therapy. **D.** Oncogenetic trajectories for diagnosis and relapse TALL42 and TALL30 samples. **E.** UMAP visualizations of Dx and Rel primary samples from TALL42 case analysed by short read scRNA sequencing. UMAP plots of 14.049 single leukemic cells coloured by cell identity (Diagnosis vs Relapse; upper panel) and cluster-based by transcriptomic data (fourteen different clusters; lower panel). The red dotted ellipse highlights cluster 6 in TALL42D sample. **F.** Dot plot visualization of biomarker genes average expression selected from the unsupervised differential expression analysis between Rel and Dx samples from TALL42 case. Average expression and proportion of expressing cells for selected biomarkers are represented in each cluster from Dx and Rel samples. The red dotted line highlights the pre-relapse cells reservoir. **G-H.** GSEA of positively and negatively enriched pathways at relapse and pre-relapse stages in TALL42 samples. **G.** Representation of selected Hallmark pathways significantly deregulated in relapse (1^st^ column Rel vs Dx; *P* adj <0.05) and pre-relapse (2^nd^ column Sub vs Dx, red dotted line; *P* adj <0.05) clusters as compared to matched diagnosis counterparts. **H.** Representation of specific gene sets from the MSigDB C1 and C2 collections significantly upregulated (in red) or downregulated (in blue) at relapse (1^st^ column Rel vs Dx; *P* adj <0.05) and pre-relapse (2^nd^ column Sub vs Dx; *P* adj <0.05) as compared to matched primary diagnosis counterparts. Color code indicates the cell proliferation, invasiveness, stemness and oxidative phosphorylation signatures. **I** Single-cell scores for several leukemia stem cell (LSC) signatures in TALL42 diagnosis (Dx), pre-relapse (Sub) and relapse (Rel) clusters. The box includes the median, hinges mark the 25^th^ and 75^th^ percentiles. *P* values from a two-sided paired t-test are shown. ns, not significant; ****, *p* < 0.0001 **J.** UMAP visualizations of Dx and Rel primary samples from TALL42 case analysed by long read single-cell RNA sequencing. UMAP plots of 14.883 single leukemic cells (8.861 cells from diagnosis and 6.022 cells from relapse) coloured by cell identity (Diagnosis vs Relapse; left panel) and cluster-based by transcriptomic data (9 different clusters corresponding to leukemic cells; right panel). **K.** Projection of *TP53* WT and mutant (*TP53*-R175H) reads at chromosomal location chr17:7’675’088 (hg38) from TALL42 Dx and Rel samples. **L.** Quantification of *TP53* mutant reads retrived in isogenic Dx (6/193) and Rel (35/35) TALL42 samples. **M.** Single-cell signature scores for relevant hallmark pathways of the relapse expression signature (OXPHOS, MYC target V1 and MTORC1 signaling) in *TP53* WT and mutant cells. The box includes the median, hinges mark the 25^th^ and 75^th^ percentiles.

Comparison of the diagnosis and relapse samples uncovered a minor cell population in two diagnosis samples (TALL42 and TALL30) that clustered nearby their paired relapse counterparts, suggesting the existence of a “pre-relapse” cells transcriptional state at diagnosis (**Supplementary Figures 4C, 4E**). We further integrated the combined analyses of mutation, copy number and TCR rearrangement to reconstitute leukemia architecture before and after treatment along with single-cell expression profiling in these two cases (**Figure 5D**).

*In case TALL42*, the small cluster of cells represented approximately 10% of the total leukemic cells at diagnosis (9.6% and 15% of the primary and PDX samples, respectively) (**Figures 5E-F** and **Supplementary Figures 4G-Q**). This minor cell population displayed both a distinct TCRB rearrangement (TCRB#2) and a distinct second-allele 9p/*CDKN2A* deletion from the dominant clone at diagnosis, and no 17p deletion, while the relapse population displayed the same subclonal TCRB#2 but yet another 9p/*CDKN2A* deletion (specific to relapse) and both the *TP53*-R175H and del17p/*TP53* (those last being not detected by NGS and inference analysis in the 10% subclone at diagnosis) (**Figure 5D** and **Supplementary Figures 4I, 4K, 4P, 4R**). Interestingly, the minor subpopulation at diagnosis was enriched in most of the relapse expression signature including OXPHOS, MTORC1 signalling, MYC targets V1, E2F targets, cholesterol homeostasis and DNA repair, these pathways being even more upregulated in the matched relapse cells (**Figure 5G** and **Supplementary Figure 4S**). We interrogated additional geneset collections and found a significant enrichment of stemness, invasiveness and OXPHOS signatures combined with downregulation of cell proliferation pathways, in line with the properties attributed to a quiescent, LIC-activity enriched, pre-relapse leukemia reservoir (**Figure 5H** and **Supplementary Figure 4T**). Specifically, interrogation of several leukemia stem cell (LSC) signatures^37–40^ showed that this cluster had increased LSC signature scores compared to diagnosis cells, as in relapse cells (**Figure 5I**).

*In the second case, TALL30*, a minor cell population representing approximately 5% at diagnosis, trackable by the same TCRB rearrangement as relapse cells but distinct from the dominant diagnostic population, also demonstrated the deregulation of the relapse signature (**Supplementary Figure 5**).

Clearly, the very low frequency of the *TP53* mutation in the two cases (VAF <0.02% as found by ddPCR) contrasted with the 5-10% frequency of the minor cell population expressing the relapse signature. In case TALL42, the clonal architecture showed that both the 5-10% minor clone population and the *TP53^alt^* mutation derived from a common ancestor bearing the same TCRB rearrangement from which they diverged by acquiring a distinct 9p/*CDKN2A* second-allele deletion in addition to the *TP53* mutation in the relapse clone (**Figure 5D**). In case TALL30, the *TP53*-mutated cells may have emerged from the minor cell population or from an ancestor pre-relapse reservoir. Although somewhat counterintuitive, these data suggest that relapse may be primed in a pre-existing reservoir population expressing the relapse expression signature, the *TP53* mutation occurring in a fraction of those cells that further expand upon treatment. To investigate this hypothesis, we performed deep-sequencing, long-read Nanopore scRNAseq on the 10X cDNAs to track the *TP53* mutations at the single-cell resolution, allowing to directly assess the transcriptional profile of the very rare *TP53*-mutant cells. In TALL42, we obtained a final mean depth of 110X at diagnosis and 40X at relapse, in 8.861 cells and 6.022 cells respectively. The expected *TP53-*R175H missense mutation was retrieved in all (35/35) covered reads from the del17p relapse sample, but also in rare (6/193) reads from the diagnosis sample (**Figures 5J-L** and **Supplementary Figures 4U-V**). While not directly numerically comparable to ddPCR since they operate at different biological and technical levels (allele level quantification versus transcriptional profiling), these data confirmed the enrichment of the relapse expression signature in the rare *TP53*-mutated cells as compared to the vast majority of *TP53 WT* cells from diagnosis (**Figure 5M**). Similar results were obtained in the second TALL30 case in which the relapse expression signature was also enriched in the rare *TP53*-mutant cells detected at diagnosis (**Supplementary Figures 5P-U**).

Collectively, our data show that *TP53*-mutated cells pre-existing at diagnosis constituted a pre-relapse reservoir population expressing the relapse signature, that expanded upon treatment along with sequential inactivation of the second *TP53* allele.

## DISCUSSION

Treatment failure and subsequent patient death for ALL are largely related to the emergence, within the first 2-years, of a relapse much less responsive to therapies than the initial diagnosis leukemia. Comprehensive longitudinal genomic studies on patient cohorts showed that rather than being subtype dependent, resistance initiates upon treatment by the selection of residual, ancestor T-ALL cells surviving therapy and re-expanding^13–19^. However, no clear genomic driver of relapse were detected in most cases and the underlying mechanisms, possibly non-genetic, remain elusive. Here, we leveraged *TP53*^alt^ T-ALL relapse cases, i.e. leukemia with a clear, genetically-trackable driver of resistance to develop faithfull *in vivo* models, based on isogenic diagnosis-relapse paired xenografts, and analysed cellular features driving cell survival and re-expansion. Contrasting with many other T-ALL cases where we previously shown that the PDX resulting from diagnosis more closely resembled relapse samples than bulk diagnosis samples^13^, diagnosis and relapse features were faithfully captured in these isogenic PDX pairs resulting from *TP53*^alt^ T-ALL. This substantial difference may be due to the *TP53* lesions nature confering an *in vivo* advantage upon treatment only, as seen in other premalignant conditions^18,41^. Importantly, these isogenic PDX models enable to overcome fundamental limitations of *ex vivo* functional studies on primary samples or cell lines, i.e. rarity and practical workability for primary samples; highly rearranged and drifted cells for cell lines; and, more generally, a lack of microenvironnement support. In addition, isogenic PDXs also allow to avoid bias related to inter-patient heterogeneity, such as the markedly distinct proliferation range between our four T-ALL cases, faithfully captured by the isogenic PDX pairs (**Supplementary Figure 1B**).

Interestingly, we found that the *TP53*^alt^-driven T-ALL relapse cells did not display strong genomic instability, suggesting a direct oncogenic role of *TP53* alterations in those leukemias as “gatekeeper” rather than its commonly recognized “caretaker” role. This was confirmed by *TP53* silencing experiments evidencing a cell-intrinsic gain of malignancy in relapse cells largely driven by the *TP53* inactivation itself. The relapse phenotype we identified includes a cell-intrinsic functional gain of LIC and proliferation, along with an expression profile that involved cell metabolism, signaling, MYC pathways, and stemness, consistently with a role of *TP53* in the regulation of normal and pathologic stem cell function in hematopoiesis^42^. We especially evidenced functionally that increased *TP53*^alt^-dependent OXPHOS was crucial to the relapse phenotype, as may be seen in other acute leukemia subtypes such as acute myeloid leukemia or various solid cancers^43,44^.

By combining single-cell transcriptomics, genomics and ultra-sensitive specific backtracking by ddPCR at diagnosis, we could reconstitute the clonal architecture at diagnosis and relapse in two cases (**Figure 5D**). We detected the expected *TP53* mutations at diagnosis at ultra-low frequencies, showing that the relapse originated from a pre-existing ultra-rare reservoir. Interestingly, a remission sample from one case could be injected and expanded towards an overt T-ALL PDX that was *TP53*-mutatedbut not yet 17p deleted, demonstrating the sequential bi-allelic acquisition through leukemia progression. Intriguingly, a transcriptional cluster representing a cell population of approximately 5-10% of cells was detected in two cases at diagnosis, although the vast majority of those cells were *TP53*^WT^. In one of these two cases (TALL42), clonal architecture demonstrated that a rare *TP53*-mutated clone and the minor (5-10%) cell population issued from the same ancestor, distinct from the dominant clone at diagnosis. In the second case, the *TP53*-mutation may have emerged directly from the minor cell population expressing the relapse signature or from an ancestor. These data raise the possibility that the *TP53* mutation would stabilize a pre-existing transcriptional state, possibly non genetically-driven. Taken the similarity of the relapse signature and the effects of *TP53* inactivation, this minor population may exhibit transcriptional features associated with the relapse because of a functional, rather than genetic, inactivation of TP53 signaling. Beyond the *TP53*-associated relapses we explored in this study, many T-ALL relapses do not display any additional “relapse-associated genes”^17^, suggesting that leukemia resistance and relapse might be driven by non-genetic mechanisms. In line with this observation, several studies have raised a spectrum of key epigenetic, transcriptionnal remodeling, cell plasticity and microenvironmental mechanisms that may contribute to relapse^17,35,37,40,48,49^. It will thus be of interest to study whether the *TP53*^alt^-driven relapse phenotype we identified here can also be relevant in T-ALL cases lacking *TP53* genetic alterations at relapse, suggesting non-genetic driver mechanisms.

Like in the vast majority of cancers, *TP53*^alt^ T-ALL relapses are associated with very poor clinical outcomes and limited therapeutic options, representing a major clinical challenge. Detecting very rare pre-existing *TP53*-mutant cells at diagnosis, i.e., below usual thresholds, potentially combined with single-cell transcriptomic detecting high-risk cellular state, and *TP53*^alt^ bi-allelic deletion tracking, may be of broad interest for patient monitoring and early intervention with targeted therapies or hematopoietic stem cell transplantation.

## METHODS

### Patient cohort, collection of biological material and annotations

Primary samples from 42 T-ALL patients who have relapsed were obtained from blood or bone marrow (BM) aspirates/biopsies at initial diagnosis, remission (non tumoral, considered as germline reference) and first relapse. The cohort included children and young adults (age, 1 to 42 years old). Samples were stored in the Tumor Bank of Saint-Louis Hospital, Paris (BiRTH, Biobank for Research in Translational Hematology) accordingly to the French law, including informed consent. Annotations of T-ALL mutations were obtained using the NGS sequencing and analysis pipeline from Saint-Louis Hospital, using a VAF threshold of 2%^50^. Seven of the 42 patients (16.7%) had *TP53* alterations at relapse, including 4 for which live cells from both diagnosis and relapse could be recovered and injected to mice. IRB approval from the Inserm was given to this study under the number #12–078.

### Animal studies

Mice experiments described in this study comply with the current ethical regulations. Authorizations for animal experimentation were obtained from the *Ministère de l’enseignement supérieur et de la recherche* with the No. APAFIS#19958-2019032615074625v2 and APAFIS#47448-2024020909144467v3. Mice were housed under a dark-light cycle at ambient temperature (≈20-24°C) with a humidity of 40-60% and handled in a pathogen-free animal facility in accordance with the guidelines of the Animal Care and Use Committee (IRSL, Université Paris Cité). Animals were excluded from experiments if any sign of stress were observed without clinical signs of full-blown leukemia development (i.e. absence of human leukemic cells in blood, bone marrow and spleen).

### Patient derived xenografts (PDXs) of isogenic diagnosis-relapse T-ALL samples

Isogenic diagnosis (Dx) and relapse (Rel) T-ALL patient samples were thawed, qualified and injected intravenously (unless otherwise stated) at equivalent cell doses into sublethally irradiated immunodeficient NOD.Cg-Prkdc*_scid_* IL2rg*_tm1Wjl_*/SzJ (NSG) mice from the Jackson Laboratory that are bred locally, as described previously^50^. Leukemia development was monitored monthly by flow cytometry (FACS) analysis of blood or BM aspirates using hCD45 and hCD7-based staining protocol detailed in the ‘Flow cytometry’ section to confirm disease engraftment.

Human T-ALL cells from both patients and derived PDXs (BM of fully leukemic mice with >95% blasts) were used for genomics, molecular and functional experiments. To evaluate the stability of the phenotype from PDX models, leukemic cells from PDXs at first passage were serially transplanted into NSG recipient mice by intravenous injection (unless otherwise stated) at equivalent cell doses. In some circumstances to evaluate post homing involvement, leukemic cells from isogenic Dx-Rel PDXs were injected into sublethally irradiated mice by intrafemoral route. Recipient mice were monitored daily by visual and clinical observations and monthly for biological parameters, i.e. signs of stress (weakness, weight loss, ruffled coat, lethargy or ataxia) and the presence of human leukemic cells in blood and/or BM. To monitor survival, mice were sacrificed at or before reaching ethical endpoints. Pathology and flow cytometry analyses confirmed full-blown T-ALL disease with marrow, spleen and liver infiltration by leukemic cells.

### Flow cytometry

Leukemic cells obtained from peripheral blood after Ficoll separation and from BM aspirates were washed and resuspended in PBS (*Gibco*) supplemented with 2% FBS (*Gibco*) before staining. Human leukemic cells infiltration in NSG recipient mice was measured in legs, spleen and liver that were crushed or mashed. Leukemic cells suspensions were washed and resuspended in PBS supplemented with 2% FBS for staining. Human leukemic cells were then stained for 30 min at 4°C with anti-human CD45 (APC or VioBlue) and CD7 (PE or PE-Vio770) antibodies (*Miltenyi Biotec*) (1:100 dilution). Cells were then washed and analysed on a FACS Canto II or LSR Fortessa analyzer (*BD Biosciences*). Flow cytometry analyses were performed using Flowjo 10.10 (*TreeStar*).

### Whole Exome Sequencing Analysis of primary and PDX T-ALL samples

Primary samples from matched tumoral (initial diagnosis and first relapse) and non-tumoral (remission) obtained from T-ALL patients and the corresponding isogenic Dx-Rel PDXs were subjected to deep whole-exome sequencing (WES). DNA libraries were prepared using SureSelect XT Target Enrichment according to the manufacturer’s instructions (*Agilent Technologies*, Santa Clara, CA, USA). Exon capture was performed with Agilent SureSelect Human All Exon V5 kit. The samples were sequenced on an Illumina NextSeq 500 platform (*Illumina*, San Diego, CA, USA) in a 2 × 100 bp paired-end sequencing run. The average sequencing depth was 110X. Raw data were first submitted to quality check using Trim_Galore (v0.6.6) to perform quality and adapter trimming for paired-end reads. The paired end clean reads were aligned to the human reference genome (hg19) using BWA (v0.7.17-r1188). Duplicates reads were flagged using Picard-tools (v2.3.0) and local realignment around InDels and base quality score recalibration was performed using Genome Analysis ToolKit (GATK v4.2.5.0).

Somatic mutations were called by VarScan somatic (v2.4.2) and annotated by ANNOVAR (version 2022-04-13) before filtering. Somatic copy number variants (CNVs) were detected by facets (v0.16.0-1) and by an in-house R script. The final retained abnormalities were validated by visual inspection considering a number of markers ≥2 and log2ratio with respect to the individual background noise.

Exon capture was performed with Agilent SureSelect Human All Exon V6 UTR kit for the isogenic Dx-Rel PDX pairs. The same procedure as above was used to analyse PDX samples (n= 8) using the corresponding primary non-tumour sample as reference for somatic calling variant step. The overlap between V5 and V6_UTR capture was performed for the upstream analysis.

### *In vivo* limiting dilution experiments

Leukemic cells from paired Dx-Rel PDXs (primografts) were intravenously injected under limiting dilution conditions from 10^6^ to 1 cell per sublethally irradiated mouse (up to 6 mice per condition). Leukemia development in recipient NSG mice was monitored monthly by blood or BM aspirates. All mice were sacrificed at or before reaching ethical endpoints. Blast infiltration was confirmed as detailed above. Pathology analysis confirmed full-blown T-ALL diagnosis, with BM, spleen and liver infiltration by lymphoblasts expressing human CD45 and CD7 markers. Leukemia initiating cell (LIC) frequencies of isogenic Dx-Rel T-ALL PDXs were calculated from the number of engrafted mice for each cell dose in each condition using the WEHI bioinformatic ELDA (Extreme Limiting Dilution Analysis) software^51^. *P* values were calculated with overall tests for differences in LIC frequencies between any of the group tested.

### In vitro long term culture-initiating cell experiment

Leukemic cells from isogenic Dx-Rel T-ALL PDX samples (primografts) and from paired control mCherry^+^ and *TP53* KD GFP^+^ PDX samples at diagnosis were sorted by flow cytometry based on CD45^+^CD7^+^ (+/-GFP^+^ or mCherry^+^) and seeded on irradiated (30 Gy) confluent stromal cells under limiting dilution conditions from 78.125 to 1 cell per well (24 wells per condition). Leukemic cells were then cocultured on stromal feeder cells for 2 to 4 weeks under hypoxic condition (3% O_2_) as previously described^50^. Cell medium was partially changed twice a week. Positive wells for leukemic cells outgrowth were visually scored at regular time intervals. Leukemic cells detection was confirmed by flow cytometry analysis of recovered wells after 2 to 4 weeks of culture upon hypoxia as detailed above to confirm the presence of leukemic cells. LIC frequencies were calculated from the number of positive wells for leukemic cells outgrowth for each cell dose in each condition using the WEHI bioinformatic ELDA software^51^. *P* values were calculated with overall tests for differences in LIC frequencies between any of the group tested.

### Real-time measurement of leukemic cells proliferation and apoptosis with the IncuCyte® Live-Cell Analysis System

Real-time measurement of *in vitro* leukemic cell proliferation and apoptosis was performed using the IncuCyte® Live-Cell Analysis System (*Sartorius*). Human CD45^+^CD7^+^ leukemic cells from isogenic Dx-Rel T-ALL PDX samples (primografts) were sorted by flow cytometry and 100.000 cells were seeded on 96-well flat bottom plate (4 wells per condition) previously coated with 0.4μg of purified human Delta4-Fc fusion protein (*PX’Therapeutics*) according to manufacturer’s recommendation. Leukemic cells were cultured as previously described^50^ for up to 96 hours under normoxia in the IncuCyte® system (5% CO_2_ and 37 °C). Real-time *in vitro* leukemic cell growth was monitored using live-cell time-lapse imaging by capturing phase contrast images every 2 hours and analysed using the integrated confluence algorithm provided. Cell proliferation was represented as the proportion of phase objects confluence over time, without labels. Leukemic cell apoptosis was detected using the IncuCyte® Caspase-3/7 green reagent for apoptosis (*Sartorius*). Briefly, leukemic cells were cultured as previously described in the presence of 5μM Caspase-3/7 green reagent. Real-time live-cell imaging of apoptosis was monitored by capturing phase contrast and green fluorescence images every 2 hours and analysed using the integrated algorithm provided. Leukemic cell apoptosis was represented as the proprotion of green fluorescently labelled nuclei after activated caspase-3/7 cleavage of inert Incucyte® Caspase-3/7 dye.

### Apoptosis and cell proliferation end-point assays

Leukemic cells from isogenic Dx-Rel PDX T-ALL samples were co-cultured on stromal cells^50^ for 48 hours under normoxia in a humidified 5% CO_2_ incubator at 37°C. To perform apoptosis assay, leukemic cells from co-cultures were then collected, washed and stained as detailed above. Leukemic cells were then fixed and stained with PE or Biotin anti-Cleaved Caspase 3 antibody (*BD Biosciences*) as recommended by the manufacturer’s recommendations, washed and then analyzed by flow cytometry on FACS Canto II or a LSR Fortessa analyzers (*BD Biosciences*). Cell cycle assays were performed using the APC BrdU Flow kit (*BD Pharmigen*) as recommended by the manufacturer’s recommendations. Briefly, leukemic cells co-cultured on stromal cells for 48 hours were pulse-labeled for one hour at 37°C with 1mM Bromodeoxyuridine (BrdU). Leukemic cells from the recovered co-cultures were stained with cell surface antibodies as above. Leukemic cells were then fixed, permeabilized, incubated with DNase and stained with APC-conjugated anti-BrdU antibody (1:100 dilution) for 20 min at room temperature and incubated with Vybrant violet (1:1000 dilution, *Invitrogen*) for 30 min at 37°C before FACS analyses.

### *In vivo* homing assay

Leukemic cells from isogenic Dx-Rel PDXs or from matched *TP53*KD and controls Dx PDXs were injected intravenously (2-6×10^6^ cells/mouse) into sublethally irradiated immunodeficient NSG recipient mice (n = 2 to 4 mice per group). Homing capacities of human leukemic cells were evaluated 24 hours after intravenous injection by FACS analysis of BM, spleen and liver.

### *TP53* gene silencing in human leukemic cells and *in vivo* transplantations

The Hu ShRNA anti-*TP53* was previously described^52^. The human *TP53* cDNA was cloned under the H1 promoter in the backbone pRRL.SIN.cPPT.PGK.GFP lentiviral vector (*Addgene* plasmid #22038). The pRRL_H1shCTL_PGK_mCherry lentivector was constructed in house from the pRRL_H1shLacZ_PGK_eGFP mentioned above. Lentiviral particles were produced as described^53^ with jetPRIME reagents (*Polyplus transfection*) by co-transfection of viral production plasmids, according to the manufacturer’s protocol. Lentiviral supernatants titration was performed using the qPCR Lentivirus Titer Kit (*Applied Biological Materials*) according to the manufacturer’s recommendation.

Leukemic cells from primary PDX samples at diagnosis were cultured on 24-wells plates previously coated with 2.5μg of purified human Delta4-Fc fusion protein (*PX’Therapeutics*) and transduced for 48-72 hours with the lentiviral supernatants including controls (at MOI 20-50) in the presence of 4ug/ml Protamine sulfate (*Sigma*). Transduced leukemic cells were then washed , GFP- and mCherry-FACS sorted and immediately intravenously injected into sub-lethally irradiated NSG mice, at equivalent dose in parallel settings. For competitive experiments, leukemic cells transduced with a shRNA CTL mCherry or shRNA *TP53*-GFP lentivirus were FACS-sorted, and mixed in equivalent number before intravenous injection into sub-lethally irradiated NSG mice. The starting GFP^+^/mCherry^+^ ratio was re-checked by flow cytometry before IV-injection. Gene silencing efficiency was checked by immunoblot (human p53 D0-1, *Santa Cruz*; Actin,*Santa Cruz*).

### Large-scale gene expression analysis of primary and PDX T-ALL samples by bulk RNA sequencing analysis

Total RNAs were extracted from frozen pellets of leukemic blasts at initial diagnosis and first relapse (n= 8 samples) and from BM of fully leukemic NSG mice for the corresponding derived PDXs (n= 24 PDX samples, first injection) using Qiagen RNA extraction kit (*Qiagen,* miRNeasy Tissue cell advance) according to the manufacturer’s recommendations. cDNA libraries were prepared using the TruSeq Stranded mRNA kit (*Illumina*, San Diego, CA, USA). Alignment to the hg19 (GRCh37) reference genome was performed using STAR aligner (Version 2.7.8a) with quantMode GeneCounts parameter to build raw count matrix for gene-level summarisation based on the in-built hg19 RefSeq-based annotation. Normalisation by the method of trimmed mean of M-values (TMM) was performed using the calcNormFactors function in edgeR. A linear model fitted in limma package was performed to identify differential gene expression between relapse and diagnosis samples in basal conditions. Differentially expressed genes were identified based on the adjusted *p*-value < 0.05 and filtered with a fold change absolute value > 0.5. The relationship between samples was visualized by hierarchical clustering and principal component analysis (PCA). Over Representation Analysis (ORA) and Gene set enrichment analyses (GSEA) were then performed to identify significant - hallmark, positional (C1), curated (C2), and ontology (C5) - genes sets using the fgsea and msigdbr packages. Additional analyses on PDX samples from secondary transplants were performed using the ClariomTM D Assay plateform (*Thermo Fisher Scientific*) using the manufacturer’s recommendations.

### Mitochondrial Respiration and Glycolytic Function Measurements

Oxygen consumption rate (OCR) and extracellular acidification rate (ECAR) were respectively measured with the Seahorse XFp Cell Mito Stress and XFp Glycolysis Stress Test kits on the Seahorse XFp Analyser (*Agilent, Seahorse Bioscience*) in accordance with the manufacturer’s recommendations. Leukemic cells recovered from 24h co-culture on stromal cells were seeded at 4 x 10^5^ cells per well (optimal cell seeding density was determined in preliminary experiments) in an XFp-well miniplate previously coated with Cell-Tak (*Corning, Fisher Scientific*) in 180µl of XF DMEM Medium (pH 7.4), supplemented with 2 mM L-glutamine (*Gibco*) (for OCR and ECAR), 1 mM fresh sodium pyruvate (*Gibco*) (for OCR only) and 10 mM D-glucose (*Sigma*) (for OCR only) on the day of the assay. For OCR measurements, sequential compound injections of Oligomycin (1µM), FCCP (1µM), Rotenone (0.5 µM) and Antimycin A (0.5µM) enabled measurements of basal respiration, ATP production, proton leak, maximal respiration, spare respiratory capacity, and non-mitochondrial respiration. For ECAR measurements, sequential compound injections of Glucose (10mM), Oligomycin (1µM) and 2-Deoxy-glucose (2-DG, 50mM) enabled measurements of glycolysis, glycolytic capacity, glycolytic reserve and non-glycolytic acidification. Key parameters of mitochondrial respiration and glycolytic functions were recorded for 90 minutes after cells were placed into the analyzer. OCR (pmol/min) and ECAR (mpH/min) were measured simultaneously in all wells, three times at each step, by optical fluorescent O2 sensors, and a minimum of three replicates were analyzed per condition in any given experiment. Unless otherwise stated, all compounds and material were obtained from *Agilent Seahorse Bioscience*.

### Cell lines and human T-ALL cells culture

HEK-293T cell line was purchased from ATCC (CRL-3216). HEK-293T cells were grown in DMEM medium (*Gibco*) supplemented with 2mM L-glutamine (*Gibco*), 1% penicillin/streptomycin (*Gibco*), and 10% heat-inactivated fetal bovine serum (FBS) (*Gibco*). Mouse MS5-Delta-like1^13,54^ and human hTERT-immortalized primary bone marrow MSCs^55^ derived stromal cells were maintained in αMEM medium (*Gibco*) supplemented with 2mM L-glutamine (*Gibco*), 1% penicillin/streptomycin (*Gibco*), 1μM Hydrocortisone (*Sigma*, for MSCs only) and 10% heat-inactivated FBS (*Gibco*). All cell lines were grown in a humidified 5% CO_2_ incubator at 37 °C and tested monthly negative for Mycoplasma using MycoAlert PLUS Mycoplasma Detection Kit (*Lonza,*) or MycoStrip Mycoplasma Detection Kit (*Invivogen*).

Primary and PDX human T-ALL leukemic cells were cocultured on stromal cells (MS5-DL1 or hMSC cells) in αMEM medium (*Gibco*) supplemented with 2mM L-glutamine (*Gibco*), 1% penicillin/streptomycin (*Gibco*), 10% heat-inactivated FBS (*Stem Cell Technologies*), 10% heat-inactivated human AB serum (*Institut de Biotechnologies Jacques Boy*), 20nM Human Recombinant Insuline (*SAFC Sigma*), 50ng/ml Human Recombinant SCF (*Miltenyi Biotec*), 10ng/ml Human Recombinant IL7 (*Miltenyi Biotec*), 20ng/ml Human Recombinant Flt3 ligand (*Miltenyi Biotec*) as already described^13,50^.

### *TP53*-mutant cells backtracking in isogenic primary and PDX T-ALL samples by droplet-digital PCR

Droplet-digital PCR (ddPCR) reactions were performed following Bio-Rad recommendation’s with 2X ddPCR SuperMix for probes No dUTP and commercially designed ddPCR Mutation Assays for each *TP53* mutation (*Bio-Rad*). Mutation assays are a mix of two fluorescent probes in competition for the annealing on the same target sequence: one specific probe for the wild-type sequence (FAM probe) and a HEX probe, specific of the mutated sequence. Two wet-lab validated ddPCR MUT FAMHEX Assay were used for detection of patient’s specific mutations (*TP53*-R175H (Assay ID dHsaMDV2010105) and *TP53*-R248Q (Assay ID dHsaMDV2010127)) and a non-wet-lab validated assay IS ddPCR MUT FAMHEX Assay was used for *TP53*-H214R mutation detection (Assay ID dHsaMDS2510824). Experiments were performed on genomic DNA from isogenic primary and PDX T-ALL samples. Several wells were loaded with DNA from each sample to test enough material. Droplets were generated using QX200 Droplet Generator (*Bio-Rad*). PCR reactions were performed on an Eppendorf Nexus Gradient thermocycler following cycling conditions after validation of optimum annealing/extension temperature: 95°C for 10 min, 40 cycles of two steps PCR 94°C for 30 sec, 55°C for 1 min with a 2°C/sec ramping, an enzyme denaturation step at 98°C for 10 min and a final hold at 4°C. FAM and HEX fluorescence’s of each droplet were then read on the QX200 Droplet reader. Visualization, analysis and VAFs calculation were performed with QXOne Manager 1.2 Standard edition software (*Bio-Rad*). Thresholds for each assay were determined aiming maximum specificity, despite sensitivity and quantification accuracy by only incorporated double-positive droplets of a well when at least a single positive-droplet was spotted to avoid potential false double-negative droplets (none as “rain droplets”) in the analysis. Non-template controls containing all reagents and the corresponding amount of carrier DNA to be equivalent to the tested samples were included in the analysis. VAFs were calculated by considering all the wells analysed.

### High-throughput short read single-cell RNA sequencing analysis of paired Dx-Rel primary and PDX T-ALL samples

Single-cell RNA-sequencing (scRNAseq) studies were performed on viable leukemic cells prepared from cryopreserved BM samples from T-ALL patients and the corresponding derived PDXs. Libraries from 10.000 leukemic cells were prepared using the Chromium Next GEM Single Cell 3’ v3.1 reagent kits (*10x Genomics*) and subjected to short read sequencing on the Illumina NovaSeq 6000 platform (*Illumina*, San Diego, CA, USA) in a 28-8-91 paired-end sequencing run (50.000 reads per cell) allowing the captured expression of ∼2000 genes per cell. FASTQ files were aligned to the human reference sequence (hg38) using CellRanger count pipeline (*10× Genomics*, v3.1.0). The output gene-barcode matrices were imported into the Seurat R package (v4.3.0) for scRNAseq data analysis and visualization. Cells with nFeature_RNA <500 or >20% mitochondrial genes content were filtered out. After removing of low-quality cells, data were normalized using the Normalized-Data function with “LogNormalize” and a scale factor of 10,000. Highly variable features were identified using the FindVariableFeatures function with selection method “vst”. Data were scaled using the ScaleData function and S.Score and G2M.Score obtained from CellCycleScoring function were regressed out. Cells were clustered using FindNeighbors (dims =1:50) with the FindClusters function (resolution =0.6) based on the shared nearest neighbour. Datasets were visualized through non-linear dimensional reduction using the Uniform Manifold Approximation and Projection (UMAP). To identify a list of differentially expressed genes, gene expression comparisons between clusters and/or samples were performed using the fast Wilcoxon rank-sum test and area under the receiver operating characteristic curve (auROC) analysis implemented in the R package Presto (v1.0.0). Gene signature scores were computed using the AddModuleScore_UCell function from the UCell R package (v2.6.2), which applies a rank-based scoring method for robust and scalable analysis. To detect large-scale copy number variations in tumor cells, we utilized the inferCNV package (v1.20.0)^56,57^. Gene expression profiles of all relapse cells, or of a defined subset representing the pre-relapse subclone, were contrasted against diagnostic cells to infer chromosomal variations in tumor cell clusters by analyzing expression intensities across genomic locations and thereby explore potential chromosomal alterations.

### 3’ single-cell long read sequencing on isogenic Dx-Rel primary T-ALL samples

Single-cell suspension was converted into barcoded single cell double strand cDNAs using the Next GEM Single Cell 3’ Kit (*10X Genomics*, V3.1), aiming for 10,000 cells following the manufacturer’s instructions. 10 ng of cDNA amplicons prepared using Next GEM Single Cell 3’Kits (*10X Genomics*, V3.1) were tagged with biotin using PCR amplification following the Oxford Nanopore Technologies (ONT) manufacturer’s instructions (SST_9198_v114_revK_29Jan2025) with custom-ordered oligos. [Btn]Fwd_3580_partial_read1_defined_for_3’_cDNA at 10 μM: 5’/5Biosg/CAGCACTTGCCTGTCGCTCTATCTTCCTACACGACGCTCTTCCGATCT-3’ Rev_PR2_partial_TSO_defined_for_3’_cDNA at 10 μM: 5’-CAGCTTTCTGTTGGTGCTGATATTGCAAGCAGTGGTATCAACGCAGAG-3’ and LongAmp Hot Start Taq 2X Master Mix (*NEB*).

Pull-down of the amplicons on M280 streptavidin beads (10 μg/μl, *Invitrogen*) was performed before a second PCR using the PCR Primers (PRM) included in the PCR Expansion kit (EXP-PCA001) and LongAmp Hot Start Taq 2X Master Mix (*NEB*). A standard Ligation Sequencing Kit V14 library preparation (SQK-LSK114) was completed to prepare the cDNA ends for sequencing on a PromethION device. 200 fmol of purified cDNA (Agencourt AMPure XP beads, *Beckman Coulter™*) from the average fragment size identified were quantified using an Agilent Fragment Analyzer and ligated. Only flow cells with at least 5,000 pores at the initial scan were used for sequencing. Prepared library was loaded onto R10.4.1 flow cells (FLO-PRO114M, *ONT*) on a PromethION P-48 device (*ONT*) (*Gustave Roussy Genomic Facility, Villejuif France*) with MinKNOW (v6.0.14) and super high-accuracy base-calling mode.

### 3’ single-cell long read ONT sequencing analysis

Real-time base calling was run using MinKNOW (v7.4.14) with the Dorado basecaller and the “Super-accurate basecalling v4.3.0” model. Alignments have been made using the single-cell (v2.0.3) workflow from epi2me-labs (https://github.com/epi2me-labs/wf-single-cell). After identification of the 10X adapters (from 3’ V3.1 kit, *10X Genomics*) and extraction of the cells barcodes and UMIs, the alignment on the human reference (GRCh38-2024-A, *10X Genomics*) was performed using minimap2 (v2.24) as detailed in the code, following parameters “-ax splice -uf --secondary=no”. Genes and transcripts assignment was performed using an in-house script from epi2me-labs similar to FLAMES.

Cell barcode by symbol count table were loaded in R (version 4.1.0). To call real cells from empty droplets, we used the emptyDrops() function from the dropletUtils package (version 1.12.2), which assesses whether the RNA content associated with a cell barcode is significantly distinct from the ambient background RNA present within each sample. Barcodes with *p*-value < 0.001 (Benjamini-Hochberg-corrected) were considered as legitimate cells for further analysis.

The count matrix was filtered to exclude genes detected in less than 5 cells, cells with less than 1000 UMIs or less than 200 detected genes, as well as cells with mitochondrial transcripts proportion >20%.The proportion of ribosomal gene counts and the proportion of mechanical stress-response gene counts were also estimated but not used to filter cells. Barcodes corresponding to doublet cells were identified and discarded using the union of two methods: scDblFinder (version 1.6.0) and scds (version 1.8.0) with its hybrid method with default parameters for both. We manually verified that the cells identified as doublets did not systematically correspond to cells in G2M phase.

Seurat (version 4.0.4) was applied for further data processing. The SCTransform normalization method (Hafemeister & Satija Genome Biol. 2019) was used to normalize, scale, select 3000 Highly Variable Genes and regress out bias factors (nCount_RNA, the number of detected genes). Person residuals from this regression were used for dimension reduction by Principal Component Analysis (PCA). The number of PCA dimensions to keep for further analysis was evaluated by assessing a range of reduced PCA spaces using 3 to 49 dimensions, with a step of 2. For each generated PCA space, Louvain clustering of cells was performed using a range of values for the resolution parameter from 0.1 to 1.2 with a step of 0.1. The optimal space was manually evaluated as the one combination of kept dimensions and clustering resolution resolving the best structure (clusters homogeneity and compacity) in a UMAP space. Additionally, we used the clustree method (version 0.4.3) to assess if the selected optimal space corresponded to a relatively stable position in the clustering results tested for these dimensions / resolution combinations.

An automatic annotation of cell types was performed by SingleR (version 1.6.1) (with fine-tuning step) and ClustifyR (version 1.5.1), using packages built-in references. It labels clusters (or cells) from a dataset based on similarity (Spearman correlation score) to a reference dataset with known labels. The labels with a correlation score greater than 0.25 for SingleR or greater than 0.35 for ClustifyR were kept. The annotation was made with CelliD (version 1.0.0) with genes signatures from panglao database. Marker genes for Louvain clusters were identified through a <U+00AB>one versus others<U+00BB> differential analysis using the Wilcoxon test through the FindAllMarkers() function (Seurat), considering only genes with a minimum log fold-change of 0.5 in at least 75% of cells from one of the groups compared, and FDR-adjusted p-values <0.05 (Benjaminin-Hochberg method). SNP were called by using bam-readcount on individual BAMs per barcodes.

## Statistical analyses

Statistical analyses were performed with GraphPad Prism (version 10.3; *GraphPad Software Inc.*) or R packages using Mann-Whitney, Anova, Wilcoxon and Kruskal-Wallis tests, as indicated in figure legends. Unless otherwise stated, the data are expressed as mean±s.d. of *n*=3 or more determinations.

For xenotransplantation analysis, Kaplan–Meier survival curves were performed and the statistical significance was determined with the Log-rank (Mantel-Cox) test. Overall survival was defined as the time from cell injection to euthanasia or death at predefined endpoints. Animals were censored if the cause of death/sacrifice was not leukemia related. Number of mice and median survival are indicated into brackets.

For limiting dilution assays, statistical analysis was performed using the WEHI bioinformatic ELDA (Extreme Limiting Dilution Analysis) software^51^). Differences were considered statistically significant at *p*<0.05 (*), *p*<0.01 (**), *p*<0.001 (***) or *p*<0.0001 (****).

## Authors’ Contributions

Conception and design of the study: SG, EC and JS

Development of methodology: SG, SQ, LH, RJ, MA and JS

Acquisition of data (provided materials and animals, acquired and managed patients, provided facilities): SG, SQ, LH, LM, MP, RK, HB, MB, VP, PF, AB, HD, NB, EC and JS

Analysis and interpretation of data (biostatistics, computational analysis, statistical analysis): SG, SQ, LH, LM, MB, RJ, MA, FS, EC and JS

Writing of the manuscript - original draft preparation, review and editing: SG, HdT, EC and JS

Grants acquisition and study supervision: SG and JS

All authors critically reviewed the manuscript.

## Disclosure of potential conflicts of interest

The authors declare no potential conflicts of interest.

## Acknowledgments

The authors thank Niclas Setterblad, Christelle Doliger, Antonio Alberdi, Sophie Duchez and Claire Maillard from the Technological core facility of the Saint-Louis Research Institut, Inserm UMR1342 / CNRS EMR8000; Nathalie Droin and Maëla Francillette from the Genomic core facility of Gustave Roussy; the ICGex NGS platform of the Curie Institut for high-throughput sequencing of *10X* cDNA librairies.

## Fundings

This work was supported by grants to Jean Soulier from the ERC St Grant Consolidator #311660, the team FRM 2022 program from the Fondation pour la Recherche Médicale #00107939, the Ligue Contre le Cancer IDF, the Fondation Leucémie Espoir, and by grants to Stéphanie Gachet from the Fondation de France, the Fondation ARC pour la Recherche sur le Cancer (PJA 20191209702), and the Paris Saint-Louis Leukemia Institute program (a grant managed by the French National Research Agency under the France 2030 program, ANR-23-IAHU-0005, Institut de la Leucémie Paris Saint-Louis). The ICGex platform was supported by the grants from the French National Research Agency, the Canceropole Ile-de-France and by the SiRIC-Curie program.

## Data-sharing statement

The data that support the findings of this study are publically available at EBI (https://www.ebi.ac.uk/biostudies) under the accession numbers BioStudies S-BSST2554 and S-BSST2808, or in the Supplementary materials. The custom code used for the analyses will be available on the published version of the article.

## Notes

### Competing Interest Statement

The authors have declared no competing interest.

### Summary of Updates

This version of the manuscript has been revised to update the following: Title and Abstract revised; Section on single-cell profilings with additional data; Figure 5 revised; Supplementary Figures 3, 4 and 5 revised; Supplementary Tables revised; Supplemental files updated.

## REFERENCES

1. Van Vlierberghe, P. & Ferrando, A. The molecular basis of T cell acute lymphoblastic leukemia. J Clin Invest 122, 3398–406 (2012).

2. Buckley, M., Yeung, D. T., White, D. L. & Eadie, L. N. T-cell acute lymphoblastic leukaemia: subtype prevalence, clinical outcome, and emerging targeted treatments. Leukemia 39, 1294–1310 (2025).

3. Weng, A. P. et al. Activating mutations of NOTCH1 in human T cell acute lymphoblastic leukemia. Science 306, 269–71 (2004).

4. Ferrando, A. A. et al. Gene expression signatures define novel oncogenic pathways in T cell acute lymphoblastic leukemia. Cancer Cell 1, 75–87 (2002).

5. Soulier, J. et al. HOXA genes are included in genetic and biologic networks defining human acute T-cell leukemia (T-ALL). Blood 106, 274–86 (2005).

6. Girardi, T., Vicente, C., Cools, J. & De Keersmaecker, K. The genetics and molecular biology of T-ALL. Blood 129, 1113–1123 (2017).

7. Pölönen, P. et al. The genomic basis of childhood T-lineage acute lymphoblastic leukaemia. Nature 632, 1082–1091 (2024).

8. Desjonqueres, A. et al. Acute lymphoblastic leukemia relapsing after first-line pediatric-inspired therapy: a retrospective GRAALL study. Blood Cancer J 6, e504 (2016).

9. Heikamp, E. B. & Pui, C. H. Next-Generation Evaluation and Treatment of Pediatric Acute Lymphoblastic Leukemia. J Pediatr 203, 14–24 e2 (2018).

10. Pagliaro, L. et al. Acute lymphoblastic leukaemia. Nat. Rev. Dis. Primer 10, 41 (2024).

11. Brady, S. W. et al. The genomic landscape of pediatric acute lymphoblastic leukemia. Nat. Genet. 54, 1376–1389 (2022).

12. Pölönen, P., Mullighan, C. G. & Teachey, D. T. Classification and risk stratification in T-lineage acute lymphoblastic leukemia. Blood 145, 1464–1474 (2025).

13. Clappier, E. et al. Clonal selection in xenografted human T cell acute lymphoblastic leukemia recapitulates gain of malignancy at relapse. J Exp Med 208, 653–61 (2011).

14. Mullighan, C. G. et al. Genomic analysis of the clonal origins of relapsed acute lymphoblastic leukemia. Science 322, 1377–80 (2008).

15. Tosello, V. et al. WT1 mutations in T-ALL. Blood 114, 1038–1045 (2009).

16. Anderson, K. et al. Genetic variegation of clonal architecture and propagating cells in leukaemia. Nature 469, 356–61 (2011).

17. Kunz, J. B. et al. Pediatric T-cell lymphoblastic leukemia evolves into relapse by clonal selection, acquisition of mutations and promoter hypomethylation. Haematologica 100, 1442–50 (2015).

18. Wong, T. N. et al. Role of TP53 mutations in the origin and evolution of therapy-related acute myeloid leukaemia. Nature 518, 552–555 (2015).

19. Oshima, K. et al. Mutational landscape, clonal evolution patterns, and role of RAS mutations in relapsed acute lymphoblastic leukemia. Proc Natl Acad Sci U A 113, 11306–11311 (2016).

20. Meyer, J. A. et al. Relapse-specific mutations in NT5C2 in childhood acute lymphoblastic leukemia. Nat. Genet. 45, 290–294 (2013).

21. Tzoneva, G. et al. Activating mutations in the NT5C2 nucleotidase gene drive chemotherapy resistance in relapsed ALL. Nat Med 19, 368–71 (2013).

22. Tzoneva, G. et al. Clonal evolution mechanisms in NT5C2 mutant-relapsed acute lymphoblastic leukaemia. Nature 553, 511–514 (2018).

23. Dieck, C. L. et al. Structure and Mechanisms of NT5C2 Mutations Driving Thiopurine Resistance in Relapsed Lymphoblastic Leukemia. Cancer Cell 34, 136–147.e6 (2018).

24. Barz, M. J. et al. Subclonal NT5C2 mutations are associated with poor outcomes after relapse of pediatric acute lymphoblastic leukemia. Blood 135, 921–933 (2020).

25. Diccianni, M. B. et al. Clinical significance of p53 mutations in relapsed T-cell acute lymphoblastic leukemia. Blood 84, 3105–12 (1994).

26. Hof, J. et al. Mutations and deletions of the TP53 gene predict nonresponse to treatment and poor outcome in first relapse of childhood acute lymphoblastic leukemia. J Clin Oncol 29, 3185–93 (2011).

27. Hof, J. et al. NOTCH1 mutation, TP53 alteration and myeloid antigen expression predict outcome heterogeneity in children with first relapse of T-cell acute lymphoblastic leukemia. Haematologica 102, e249–e252 (2017).

28. Richter-Pechańska, P. et al. Identification of a genetically defined ultra-high-risk group in relapsed pediatric T-lymphoblastic leukemia. Blood Cancer J. 7, e523 (2017).

29. Simonin, M. et al. Prognostic value and oncogenic landscape of TP53 alterations in adult and pediatric T-ALL. Blood 141, 1353–1358 (2023).

30. Kempter, T. et al. Subclonal TP53 and KRAS variants combined with poor treatment response identify ultrahigh-risk pediatric patients with T-ALL. Blood Adv. 9, 1267–1279 (2025).

31. Li, B. et al. Therapy-induced mutations drive the genomic landscape of relapsed acute lymphoblastic leukemia. Blood 10.1182/blood.2019002220 (2020) doi:10.1182/blood.2019002220.

32. Armstrong, F. et al. NOTCH is a key regulator of human T-cell acute leukemia initiating cell activity. Blood 113, 1730–1740 (2009).

33. Liu, Y., Su, Z., Tavana, O. & Gu, W. Understanding the complexity of p53 in a new era of tumor suppression. Cancer Cell 42, 946–967 (2024).

34. Dong, Y., Tu, R., Liu, H. & Qing, G. Regulation of cancer cell metabolism: oncogenic MYC in the driver’s seat. Signal Transduct. Target. Ther. 5, 124 (2020).

35. Richter-Pechańska, P. et al. Pediatric T-ALL type-1 and type-2 relapses develop along distinct pathways of clonal evolution. Leukemia 36, 1759–1768 (2022).

36. Albertí-Servera, L. et al. Single-cell DNA amplicon sequencing reveals clonal heterogeneity and evolution in T-cell acute lymphoblastic leukemia. Blood 137, 801–811 (2021).

37. Calvo, J. et al. High CD44 expression and enhanced E-selectin binding identified as biomarkers of chemoresistant leukemic cells in human T-ALL. Leukemia 39, 323–336 (2025).

38. Xu, J. et al. A multiomic atlas identifies a treatment-resistant, bone marrow progenitor-like cell population in T cell acute lymphoblastic leukemia. *Nat*. Cancer 6, 102–122 (2025).

39. Ebinger, S. et al. Characterization of Rare, Dormant, and Therapy-Resistant Cells in Acute Lymphoblastic Leukemia. Cancer Cell 30, 849–862 (2016).

40. Costea, J. et al. Role of stem-like cells in chemotherapy resistance and relapse in pediatric T-cell acute lymphoblastic leukemia. Nat. Commun. 16, 5413 (2025).

41. Hsu, J. I. et al. PPM1D Mutations Drive Clonal Hematopoiesis in Response to Cytotoxic Chemotherapy. Cell Stem Cell 23, 700–713.e6 (2018).

42. Chen, S. et al. Mutant p53 drives clonal hematopoiesis through modulating epigenetic pathway. Nat. Commun. 10, 5649 (2019).

43. Liu, Y., Su, Z., Tavana, O. & Gu, W. Understanding the complexity of p53 in a new era of tumor suppression. Cancer Cell 42, 946–967 (2024).

44. Chomczyk, M. et al. Impact of p53-associated acute myeloid leukemia hallmarks on metabolism and the immune environment. Front. Pharmacol. 15, 1409210 (2024).

45. Chiu, P. P. L., Jiang, H. & Dick, J. E. Leukemia-initiating cells in human T-lymphoblastic leukemia exhibit glucocorticoid resistance. Blood 116, 5268–5279 (2010).

46. Belmonte, M., Hoofd, C., Weng, A. P. & Giambra, V. Targeting leukemia stem cells: which pathways drive self-renewal activity in T-cell acute lymphoblastic leukemia? Curr Oncol 23, 34–41 (2016).

47. Gerby, B. et al. High-throughput screening in niche-based assay identifies compounds to target preleukemic stem cells. J. Clin. Invest. 126, 4569–4584 (2016).

48. Yadav, B. D. et al. Heterogeneity in mechanisms of emergent resistance in pediatric T-cell acute lymphoblastic leukemia. Oncotarget 7, 58728–42 (2016).

49. Ariës, I. M. et al. PRC2 loss induces chemoresistance by repressing apoptosis in T cell acute lymphoblastic leukemia. J. Exp. Med. 215, 3094–3114 (2018).

50. Gachet, S. et al. Deletion 6q Drives T-cell Leukemia Progression by Ribosome Modulation. Cancer Discov 8, 1614–1631 (2018).

51. Hu, Y. & Smyth, G. K. ELDA: extreme limiting dilution analysis for comparing depleted and enriched populations in stem cell and other assays. J. Immunol. Methods 347, 70–78 (2009).

52. Ceccaldi, R. et al. Bone marrow failure in Fanconi anemia is triggered by an exacerbated p53/p21 DNA damage response that impairs hematopoietic stem and progenitor cells. Cell Stem Cell 11, 36–49 (2012).

53. Sebert, M. et al. Clonal hematopoiesis driven by chromosome 1q/MDM4 trisomy defines a canonical route toward leukemia in Fanconi anemia. Cell Stem Cell 30, 153–170.e9 (2023).

54. Gerby, B. et al. Optimized gene transfer into human primary leukemic T cell with NOD-SCID/leukemia-initiating cell activity. Leukemia 24, 646–9 (2010).

55. Mihara, K. et al. Development and functional characterization of human bone marrow mesenchymal cells immortalized by enforced expression of telomerase. Br. J. Haematol. 120, 846–849 (2003).

56. Patel, A. P. et al. Single-cell RNA-seq highlights intratumoral heterogeneity in primary glioblastoma. Science 344, 1396–1401 (2014).

57. Zhu, Q. et al. Single cell multi-omics reveal intra-cell-line heterogeneity across human cancer cell lines. Nat. Commun. 14, 8170 (2023).

