## Supplementary material for "*In vivo* isogenic modelling unveils *TP53*-associated relapse trajectories in T-cell acute lymphoblastic leukemia"

**This document includes:**

Supplementary Figures 1 – 5

Supplementary Tables 3

**The following supplementary files are available as separate .xls files:**

Supplementary Tables 1 – 2 – 3 (Excel files)

**List of Supplementary items:**

**Supplementary Figure 1:** *TP53*$^{alt}$ T-ALL relapse cells display a cell-intrinsic gain of malignancy phenotype.

**Supplementary Figure 2:** *TP53* silencing in diagnosis PDX samples recapitulates the relapse phenotype.

**Supplementary Figure 3:** TP53$^{alt}$ T-ALL relapses are characterized by deregulated cell metabolism.

**Supplementary Figure 4:** Single-cell profiling of isogenic Dx-Rel T-ALL samples unveils pre-relapse subclones at diagnosis.

**Supplementary Figure 5.** Single-cell profiling of Dx-Rel pairs from TALL30 case.

**Supplementary Table 1:** Somatic mutations details from WES analysis on paired diagnosis-relapse primary and PDX T-ALL samples.

**Supplementary Table 2:** CNVs details from WES analysis on paired diagnosis-relapse primary and PDX T-ALL samples.

**Supplementary Table 3:** Limit dilution assays (LDA) details from *in vivo* and *in vitro* experiments.

#### SUPPLEMENTARY FIGURE LEGENDS

**Supplementary Figure 1. TP53$^{alt}$ T-ALL relapse cells display a cell-intrinsic gain of malignancy phenotype. A.** Engraftment follow up of primary human leukemic cells in the bone marrow (BM) of NSG mice injected with equivalent cell doses of isogenic diagnosis (Dx) and relapse (Rel) samples from 4 T-ALL patients (right to left panels: TALL11, TALL30, TALL42, and TALL43 cases). **B.** Kaplan-Meier survival curves of NSG mice engrafted with isogenic Dx-Rel primary samples from the 4 T-ALL patients depicted in panel A. **C-F.** Secondary (C-D) and tertiary injections (E-F) of isogenic Dx-Rel PDXs established from the 4 T-ALL cases depicted in panels A and B. **C.** Engraftment follow up of human leukemic cells in BM of NSG mice injected with 3 cell doses ($10^6$, $10^5$, $10^4$) of isogenic Dx-Rel PDX samples (primografts) from 4 T-ALL cases (right to left panels: PDX11, PDX30, PDX42 and PDX43 cases). **D.** Kaplan-Meier survival curves of NSG mice engrafted with 3 cell doses ($10^6$, $10^5$, $10^4$) of isogenic Dx-Rel PDX samples (primografts) from 4 T-ALL cases (right to left panels: PDX11, PDX30, PDX42 and PDX43 cases). **E.** Engraftment of human leukemic cells in BM of NSG mice injected with 3 cell doses ($10^6$, $10^5$, $10^4$) of isogenic Dx-Rel samples established from PDX11 (2$^{nd}$ injection). **F.** Kaplan-Meier survival curves of NSG mice engrafted with 3 cell doses ($10^6$, $10^5$, $10^4$) of isogenic Dx-Rel PDX samples (2$^{nd}$ injection) from 2 T-ALL cases (right to left panels: PDX11 and PDX42 cases). **G.** LIC frequencies of isogenic Dx-Rel PDX sample pairs calculated from *in vitro* long-term culture-initiating cell assays. **H.** *In vitro* cell cycle analysis after BrdU pulse labelling of isogenic Dx-Rel T-ALL PDX samples co-cultured on stromal cells for 2 days. Statistical analyses compared Dx and Rel PDX samples of each isogenic pair for each G1, S and G2-M phases **I.** *In vitro* cell viability analysis after cleaved caspase 3 staining of isogenic Dx-Rel T-ALL PDX samples co-cultured on stromal cells for 2 days. **J-K.** *In vivo* homing assays of 4 isogenic Dx-Rel T-ALL PDX pairs. Proportions (**J**) and absolute numbers (**K**) of homed human CD45$^+$CD7$^+$ leukemic cells in BM (upper panels), spleen (middle panels) and liver (lower panels) of recipient mice (n=2 to 4). **L.** Engraftment follow up of paired Dx-Rel samples from PDX11 after intra femoral injection, in comparison to panel A. **M.** Kaplan-Meier survival curves of mice injected intrafemorally with paired Dx-Rel samples from PDX11, in comparison to panel B.

Statistical data in this figure are represented as mean ± s.d. Significance is indicated by P values; A, C, E, J-L: Mann-Whitney and Anova tests; B, D, F, M: Log-rank (Mantel-Cox) test; G: overall tests for differences using ELDA software. Number of recipient mice and median survival are indicated into brackets on Kaplan-Meier survival curves. ns: non-significant, *, $p < 0.05$; **, $p < 0.01$; ***, $p < 0.001$; ****, $p < 0.0001$.

**Supplementary Figure 2. *TP53* silencing in diagnosis PDX samples recapitulates the relapse phenotype. A.** Western blot showing *TP53* silencing efficiency in unsorted blasts from diagnosis PDX samples (PDX30D and PDX42D). **B.** Engraftment kinetics of paired control mCherry$^+$ and *TP53* KD GFP$^+$ leukemic cells from PDX11D, PDX42D and PDX43D (left, middle and right panel respectively). **C.** Kaplan-Meier survival curves of NSG mice injected in parallel with equivalent cell doses of paired control mCherry$^+$ or *TP53* KD GFP$^+$ leukemic cells from PDX11D, PDX42D and PDX43D (left, middle and right panels, respectively). **D-E.** Competitive primary transplantations of paired control mCherry$^+$ and *TP53* KD GFP$^+$ leukemic cells from PDX11D and PDX43D. Engraftment kinetics in the BM of recipient NSG mice injected with cells of both control mCherry$^+$ and *TP53* KD GFP$^+$ from PDX11D (n=4) (**D**) and PDX43D (n=5) (**E**) in competitive settings (ratio 1:1). Data are represented as ratios of *TP53* KD GFP$^+$ over control mCherry$^+$ leukemic cells within BM of the same recipient mouse and compared to the input over time. **F-H.** Secondary transplant of paired control mCherry$^+$ and *TP53* KD GFP$^+$ leukemic cells from PDX42D injected either in parallel (F-G) or in competitive (H) settings. **F.** Leukemic cells engraftment in the BM of recipient NSG mice injected with control mCherry$^+$ or *TP53* KD GFP$^+$ cells. **G.** Kaplan-Meier survival curves of NSG mice injected with sorted mCherry$^+$ or *TP53* KD GFP$^+$ leukemic cells from PDX42D. **H.** Leukemic cells engraftment in the BM of recipient NSG mice (n=4) injected with both control mCherry$^+$ and *TP53* KD GFP$^+$ cells in competitive settings (ratio 1:1). Data are represented as ratios of *TP53* KD GFP$^+$ over control mCherry$^+$ leukemic cells within BM of the same recipient mouse and compared to the input. **I.** *In vitro* cell cycle analysis after BrdU pulse labelling of paired control mCherry$^+$ and *TP53* KD GFP$^+$ leukemic cells from PDX30D co-cultured on stromal cells for 2 days. Proportions of leukemic mCherry$^+$ or GFP$^+$CD45$^+$CD7$^+$ cells in each phase of the cell cycle are indicated. Statistical analyses compared Control mCherry$^+$ to *TP53* KD GFP$^+$ samples in each G1, S and G2-M phases. **J-K.** *In vivo* homing capacities of paired control mCherry$^+$ and *TP53* KD GFP$^+$ leukemic cells from PDX42D. Proportions (**J**) and absolute numbers (**K**) of homed mCherry$^+$ and GFP$^+$ CD45$^+$CD7$^+$ leukemic cells retrieved in the bone marrow of recipient mice (n= 3 to 4) were quantified by flow cytometry. **L.** Engraftment follow up of paired control mCherry$^+$ and *TP53* KD GFP$^+$

leukemic cells from PDX30D samples after intra femoral injection, in comparison to Figure 3B. **M.** Kaplan-Meier survival curves of mice injected intrafemorally with control and *TP53* KD sorted leukemic cells from PDX30D, in comparison to Figure 3C. Statistical data in this figure, are represented as mean ± s.d. Significance is indicated by P values; B, F, J, K, L: Mann-Whitney test; D, E, H: Kruskal-Wallis test; C, G and M: Log-rank (Mantel-Cox) test. Number of recipient mice and median survival are indicated into brackets on Kaplan-Meier survival curves. ns: non-significant, *, $p < 0.05$; **, $p < 0.01$; ***, $p < 0.001$; ****, $p < 0.0001$.

**Supplementary Figure 3. *TP53*$^{alt}$ T-ALL relapses are characterized by deregulated cell metabolism. A-C.** Transcriptomic profiling of PDX samples (primografts) at diagnosis and relapse using bulk RNA sequencing. **A.** Volcano plot of differentially expressed genes between all PDXs at relapse (Rel) and their isogenic diagnosis (Dx) counterparts. Fold change (FC; x-axis) is plotted against statistical significance (adj *P* value; y-axis) for each gene. 181 genes are downregulated (FC < -0.5; $P < 0.05$) while 91 genes are upregulated (FC > 0.5; $P < 0.05$) in Rel as compared to Dx PDX samples and highlighted in blue and red respectively. *TP53* (spotted as a green dot) was significantly downregulated in relapse PDX samples. **B.** Box plots of *TP53* gene abundance (in TPM units) in RNA obtained from leukemic cells of isogenic Dx-Rel PDX pairs raised from 4 T-ALL cases (n=3 per condition). **C.** GSEA of positively and negatively enriched pathways in PDX samples from T-ALL relapses. Representation of specific gene sets from the C1 and C2 collections of the Human Molecular Signatures Database (MSigDB) significantly upregulated (in red) or downregulated (in blue) at relapse (*P* adj <0.05) in all PDX pairs pooled together (1st column "AllPDXs") and in each individual PDX pair (2nd to 5th columns; respectively PDX11, PDX30, PDX42 and PDX43). Selected signatures linked to *TP53*, cell proliferation, invasiveness, stemness and oxidative phosphorylation are depicted according to the colour code in the legend. **D-G.** Transcriptomic profiling of PDX samples after secondary transplant (2nd passage) at diagnosis and relapse using Clariom D assays. **D.** Volcano plot of differentially expressed genes between all PDXs at relapse (Rel) and their isogenic diagnosis (Dx) counterparts raised from 4 isogenic PDX pairs (n= 3 per condition). Fold change (FC; x-axis) is plotted against statistical significance (adj *P* value; y-axis) for each gene. 233 genes are downregulated (FC < -0.5; $P < 0.05$) while 283 genes are upregulated (FC > 0.5; $P < 0.05$) in Rel as compared to Dx PDX samples and highlighted in blue and red respectively. *TP53* (spotted as a green dot) was significantly downregulated in relapse PDX samples. **E.** Box plots of *TP53* gene abundance (in TPM units) in RNA obtained from leukemic cells of isogenic Dx-Rel PDX pairs raised from 4 isogenic PDX pairs at P1 (n = 3 PDX per condition). **F.** Supervised hierarchical clustering of the top fifty most differentially expressed genes in paired Dx-Rel PDX samples. Heatmap representation of the most variables genes in 4 isogenic PDX pairs. *TP53* expression was downregulated in relapse PDX samples and highlighted in red. Independent triplicates were analysed for each PDX pair. **G.** GSEA of differentially expressed genes in Rel and Dx PDX samples. Representation of selected Hallmark pathways significantly upregulated at relapse (*P* adj <0.05) in all isogenic PDX pairs pooled together (1st column "AllPDXs") and in each individual PDX pair (2nd to 5th columns; respectively PDX11, PDX30, PDX42 and PDX43). **H-I.** Key parameters evaluation of mitochondrial respiration in isogenic Dx-Rel T-ALL PDXs by real-time measurement of oxygen consumption rate (OCR) using Seahorse assays. Quantification of basal respiration (**H**) and mitochondrial spare capacity (difference between maximal and basal OCR) (**I**) in 4 independent isogenic Dx-Rel PDX pairs evaluated in 2 independent experiments. **J-K.** Direct evaluation of glycolysis in isogenic Dx-Rel T-ALL PDXs by real-time measurement of extra cellular acidification rate (ECAR) using Seahorse assays. **J.** Real-time measurements of ECAR in leukemic cells from 4 independent isogenic Dx-Rel PDX pairs (from left to right panels PDX11, PDX30, PDX42 and PDX43, respectively). **K.** Quantification of maximal glycolysis in isogenic Dx-Rel leukemic cells from 4 independent PDX pairs evaluated in 2 independent experiments. **L-M.** Key parameters evaluation of mitochondrial respiration in *TP53* KD (sh *TP53* GFP) and control (sh CTL mCherry) diagnosis PDXs using Seahorse assays. Quantification of basal respiration (**L**) and mitochondrial spare capacity (**M**) in *TP53* KD and control leukemic cells raised from 3 independent PDX diagnosis (PDXD30, PDXD42 and PDXD43) evaluated in 2 independent experiments. **N.** Quantification of maximal glycolysis in *TP53* KD and control leukemic cells raised from 3 independent PDX diagnosis (PDXD30, PDXD42 and PDXD43) evaluated in 2 independent experiments.

Statistical data in this figure are represented as mean ± s.d. from triplicates. Significance is indicated by P values; H-I and K-N: Mann Whitney test. ns: non-significant, *, $p < 0.05$; **, $p < 0.01$, ***, $p < 0.001$. ****, $p < 0.0001$.

**Supplementary Figure 4. Single-cell profiling of isogenic Dx-Rel T-ALL samples unveils pre-relapse subclones at diagnosis. A.** Bar plots representing the repartition of 117119 single leukemic cells from the sixteen samples (8 primary and 8 corresponding PDXs samples) raised from 4 T-ALL patients at Diagnosis (left) and Relapse (right). **B**. UMAP plots of 117119 leukemic cells from the sixteen samples (isogenic Dx-Rel primary and PDX samples) coloured by cell cycle phases such as G1 in pink, G2M in green and S phase in light blue. **C.** UMAP plots of 65.015 single leukemic cells from T-ALL patient samples (TALL11, TALL30, TALL42 and TALL43) at Diagnosis and Relapse coloured by samples (left panel) and disease stages (right panel; Diagnosis (green) and Relapse (purple)). The black arrow highlights a "pre-relapse" cells reservoir in TALL30D close to the TALL30R sample. **D.** Volcano plot of differentially expressed genes between TALL relapse and diagnosis samples obtained from the Wilcoxauc R command. Fold change (FC; x-axis) is plotted against statistical significance (adj *P* value; y-axis) for each gene. 207 genes are downregulated (FC < -0.5; *P* < 0.05) while 597 genes are upregulated (FC > 0.5; *P* < 0.05) in TALL relapse as compared to diagnosis samples and highlighted in blue and red respectively. *TP53* is spotted as a green dot. **E.** UMAP plots of 52.104 single leukemic from isogenic Dx-Rel PDX samples (PDX11, PDX30, PDX42 and PDX43) coloured by PDX samples (left panel) and disease stages (right panel; Diagnosis (green) and Relapse (purple)). Black arrows highlight a "pre-relapse" cells reservoir in PDX30D and PDX42D samples **F.** Volcano plot of differentially expressed genes between PDXR and PDXD samples obtained from the Wilcoxauc R command. Fold change (FC; x-axis) is plotted against statistical significance (adj *P* value; y-axis) for each gene. 136 genes are downregulated (FC < -0.5; *P* < 0.05) while 399 genes are upregulated (FC > 0.5; *P* < 0.05) in PDXR as compared to PDXD samples and highlighted in blue and red respectively. *TP53* is spotted as a green dot. **G**. Bar plots representing the repartition of 14049 single leukemic cells from TALL42 patient samples represented on UMAP plot Figure 5E (upper panel). Samples are coloured by cluster-based transcriptomic data as fourteen different clusters at Diagnosis (left) and Relapse (right). **H.** Scatter plot showing the comparative analysis of the percentage of cells expressing each gene between primary TALL42R (y axis; 'pct_out') and matched TALL42D (x axis; 'pct_in') from the Wilcoxauc R command ($\alpha$=0.05 and FC=0.15). This allows to identify biomarker genes differentially expressed between relapse and diagnosis samples. *TP53* is spotted as a green dot. **I.** UMAP visualizations of *TRBC1* (left panel) and

*TRBC2* (right panel) gene expression level in primary samples from TALL42 case at diagnosis and relapse. The red dotted ellipse highlights cluster 6 in TALL42D sample. **J.** UMAP visualization of 14049 leukemic cells from isogenic Dx-Rel TALL42 patient samples coloured by cell cycle phases such as G1 in pink, G2M in green and S phase in light blue. **K.** CNV inference analysis of cluster 6 ('Sub'; "pre-relapse" subpopulation) from TALL42D and all the clusters from TALL42R (clusters 0, 3, 4 and 9; 'Rel') interrogating copy number alterations on each chromosome (lower part) as compared to the remaining diagnosis sample (clusters 1, 2, 5, 7, 10 and 11; 'Dx') as a reference (upper part). The red circle highlights the CNV status for chromosome 17p in the pre-relapse clone (cluster 6) from TALL42D**.** **L.** UMAP visualizations of isogenic Dx-Rel PDX42 samples analysed by short read single-cell RNA sequencing. UMAP plots of 13435 single leukemic cells coloured by cell identity (Diagnosis vs Relapse; upper panel) and cluster-based by transcriptomic data (fifteen different clusters; lower panel). The red dotted ellipse highlights clusters 7 and 9 from PDX42D sample considered as the "pre-relapse" cells reservoir. **M.** Bar plots representing the repartition of 13435 single leukemic cells from PDX42 samples represented on panel L. Samples are coloured by cluster-based transcriptomic data as fifteen different clusters at Diagnosis (left) and Relapse (right). **N.** Scatter plot showing the comparative analysis of the percentage of cells expressing each gene between PDX42R (y axis; 'pct_out') and matched PDX42D (x axis; 'pct_in') from the Wilcoxauc R command ($\alpha$=0.05 and FC=0.15). This allows to identify biomarker genes differentially expressed between relapse and diagnosis samples. *TP53* is spotted as a green dot. **O.** Dot plot visualization of biomarker genes average expression selected from the unsupervised differential expression analysis between relapse and diagnosis from PDX42 samples. Average expression and proportion of expressing cells for selected biomarkers were represented in each cluster from diagnosis and relapse samples. The red dotted line highlights clusters 7 and 9 from PDX42D. **P.** UMAP visualizations of *TRBC1* (left panel) and *TRBC2* (right panel) gene expression level in isogenic Dx-Rel PDX42 samples. The red dotted ellipse highlights clusters 7 and 9 in PDX42D sample. **Q.** UMAP visualization of 13435 leukemic cells from isogenic Dx-Rel PDX42 samples coloured by cell cycle phases such as G1 in pink, G2M in green and S phase in light blue. **R.** CNV inference analysis of clusters 7 and 9 ('Sub'; "pre-relapse" subpopulation) from PDX42D and all the clusters from PDX42R (clusters 1, 10, 2, 3 and 5; 'Rel') interrogating copy number alterations on each chromosome (lower part) as compared to the remaining diagnosis sample (clusters 0, 4, 6, 8 and 11; ‘Dx) as a reference (upper part). The red circle highlights the CNV status for chromosome 17p of clusters 7 and 9 from PDX42D. **S-T.** GSEA of positively and negatively enriched pathways at relapse and pre-relapse stages in PDX42 samples. **S.** Representation of selected Hallmark pathways significantly deregulated in relapse (1$^{st}$ column Rel vs Dx; *P* adj <0.05) and “pre-relapse” (2$^{nd}$ column Sub vs Dx, red dotted line; *P* adj <0.05) clusters as compared to matched diagnosis counterparts. **T.** Representation of specific gene sets from the C1 and C2 collections that are significantly upregulated (in red) or downregulated (in blue) at relapse (1$^{st}$ column Rel vs Dx; *P* adj <0.05) and pre-relapse (2$^{nd}$ column Sub vs Dx; *P* adj <0.05) as compared to matched PDX diagnosis counterparts. Signatures linked to cell proliferation, invasiveness, stemness and oxidative phosphorylation are indicated according to the colour code. **U.** Bar plots showing the repartition of 14883 single leukemic cells from TALL42 patient samples represented on UMAP plot Figure 5J. Samples are coloured by cluster-based transcriptomic data as eleven different clusters (including leukemic and normal cells) from Diagnosis (left) and Relapse (right). **V.** UMAP visualization of *TP53* gene expression level in diagnosis and relapse primary samples from TALL42 case.

**Supplementary Figure 5. Single-cell profiling of isogenic Dx-Rel pairs from TALL30 case. A.** UMAP visualizations of diagnosis and relapse primary samples from patient TALL30 analysed by short read single-cell RNA sequencing. UMAP plots of 18997 single leukemic cells coloured by cell identity (Diagnosis vs Relapse; upper panel) and cluster-based by transcriptomic data (thirteen different clusters; lower panel). The red dotted ellipse highlights cluster 9 in primary diagnosis sample (TALL30D). **B**. Bar plots representing the repartition of 18997 single leukemic cells from TALL30 patient samples represented on UMAP plot (panel A). Samples are coloured by cluster-based transcriptomic data as thirteen different clusters at Diagnosis (left) and Relapse (right). **C.** Scatter plot showing the comparative analysis of the percentage of cells expressing each gene between primary TALL30R (y axis; ‘pct_out’) and matched TALL30D (x axis; ‘pct_in’) from the Wilcoxauc R command ($\alpha$=0.05 and FC=0.15). This allows to identify biomarker genes differentially expressed between relapse and diagnosis samples. *TP53* is spotted as a green dot. **D.** Dot plot visualization of biomarker genes average expression selected from the unsupervised differential expression analysis between relapse and diagnosis samples from TALL30 (left part) and PDX30 (right part) cases. Average expression and proportion of expressing cells for selected biomarkers were represented in each cluster from diagnosis and relapse samples. The red dotted line highlights clusters 9 and 11 from TALL30D and PDX30D, respectively. **E.** UMAP visualizations of *TRBC1* (left panel) and *TRBC2* (right panel) gene expression level in isogenic Dx-Rel TALL30 samples. The red dotted ellipse highlights cluster 9 in diagnosis sample (TALL30D). **F.** UMAP visualization of 18997 leukemic cells from isogenic Dx-Rel TALL30 patient samples coloured by cell cycle phases such as G1 in pink, G2M in green and S phase in light blue. **G.** CNV inference analysis of cluster 9 ('Sub'; pre-relapse subpopulation) from TALL30D and all the clusters from TALL30R (clusters 0, 3, 5 and 8; 'Rel') interrogating copy number alterations on each chromosome (lower part) as compared to the remaining diagnosis sample (clusters 1, 2, 4, 6, 7 and 13; 'Dx') as a reference (upper part). The red circle highlights the CNV status for chromosome 17p of cluster 9 from TALL30D. **H.** GSEA of positively and negatively enriched pathways at relapse and "pre-relapse" in primary TALL30 samples. Representation of selected Hallmark pathways significantly deregulated in relapse ($1^{st}$ column Rel vs Dx; $P$ adj <0.05) and pre-relapse ($2^{nd}$ column Sub vs Dx, red dotted line; $P$ adj <0.05) clusters as compared to matched diagnosis counterparts. **I.** UMAP visualizations of isogenic Dx-Rel PDX30 samples analysed by short read single-cell RNA sequencing. UMAP plots of 9643 single leukemic cells coloured by cell identity (Diagnosis vs Relapse; upper panel) and cluster-based by transcriptomic data (twelve different clusters; lower panel). The red dotted ellipse highlights clusters 11 in PDX30 diagnosis sample (PDX30D) considered as the pre-relapse population. **J.** Bar plots representing the repartition of 9643 single leukemic cells from PDX30 samples represented on UMAP plot (panel I). Samples are coloured by cluster-based transcriptomic data as twelve different clusters at Diagnosis (left) and Relapse (right). **K.** Scatter plot showing the comparative analysis of the percentage of cells expressing each gene between PDX30R (y axis; 'pct_out') and matched PDX30D (x axis; 'pct_in') from the Wilcoxauc R command ($\alpha$=0.05 and FC=0.15). This allows to identify biomarker genes differentially expressed between relapse and diagnosis samples. *TP53* is spotted as a green dot. **L.** UMAP visualizations of *TRBC1* (left panel) and *TRBC2* (right panel) gene expression level in isogenic Dx-Rel PDX30 samples. The red dotted ellipse highlights cluster 11 in diagnosis sample (PDX30D). **M.** UMAP visualization of 9643 leukemic cells from isogenic Dx-Rel PDX30 samples coloured by cell cycle phases such as G1 in pink,

G2M in green and S phase in light blue. **N.** CNV inference analysis of cluster 11 ('Sub'; pre-relapse subpopulation) from PDX30D and all the clusters from PDX30R (clusters 0, 2, 3, 5, 6, 7 and 8; 'Rel') interrogating copy number alterations on each chromosome (lower part) as compared to the remaining diagnosis sample (clusters 1, 4, 9 and 10; 'Dx') as a reference (upper part). The red circle highlights the CNV status for chromosome 17p of cluster 11 from PDX30D. **O.** GSEA of positively and negatively enriched pathways at relapse and pre-relapse in PDX30 samples. Representation of selected Hallmark pathways significantly deregulated in relapse (1$^{st}$ column Rel vs Dx; *P* adj <0.05) and pre-relapse (2$^{nd}$ column Sub vs Dx, red dotted line; *P* adj <0.05) clusters as compared to matched diagnosis counterparts. **P.** UMAP visualizations of Dx and Rel primary samples from TALL30 case analysed by long read single-cell RNA sequencing. UMAP plots of 17602 single leukemic cells (9721 cells from diagnosis and 7881 cells from relapse) coloured by cell identity (Diagnosis vs Relapse; left panel) and cluster-based by transcriptomic data (ten different clusters corresponding to leukemic cells; right panel). **Q.** Bar plots showing the repartition of 17602 single leukemic cells from TALL30 patient samples represented on the UMAP plot. Samples are coloured by cluster-based transcriptomic data as ten different clusters (including leukemic and normal cells) from Diagnosis (right) and Relapse (left). **R.** UMAP visualization of *TP53* gene expression level in diagnosis and relapse primary samples from TALL30 case. **S.** Projection of *TP53* WT and mutant (*TP53*-H214R) reads at chromosomal location chr17:7'674'890 (hg38) from TALL30 Dx and Rel samples. **T.** Quantification of *TP53* wt (68/72) and mutant (8/502) reads retrieved in isogenic Dx and Rel TALL30 samples. **U.** Single-cell signature scores of TP53 WT and mutant cells for relevant hallmark pathways of the relapse expression signature (OXPHOS, MYC target V1 and MTORC1 signaling) in TALL30 Dx and Rel samples. The box includes the median, hinges mark the 25$^{th}$ and 75th percentiles.

### Supplementary Figure 1

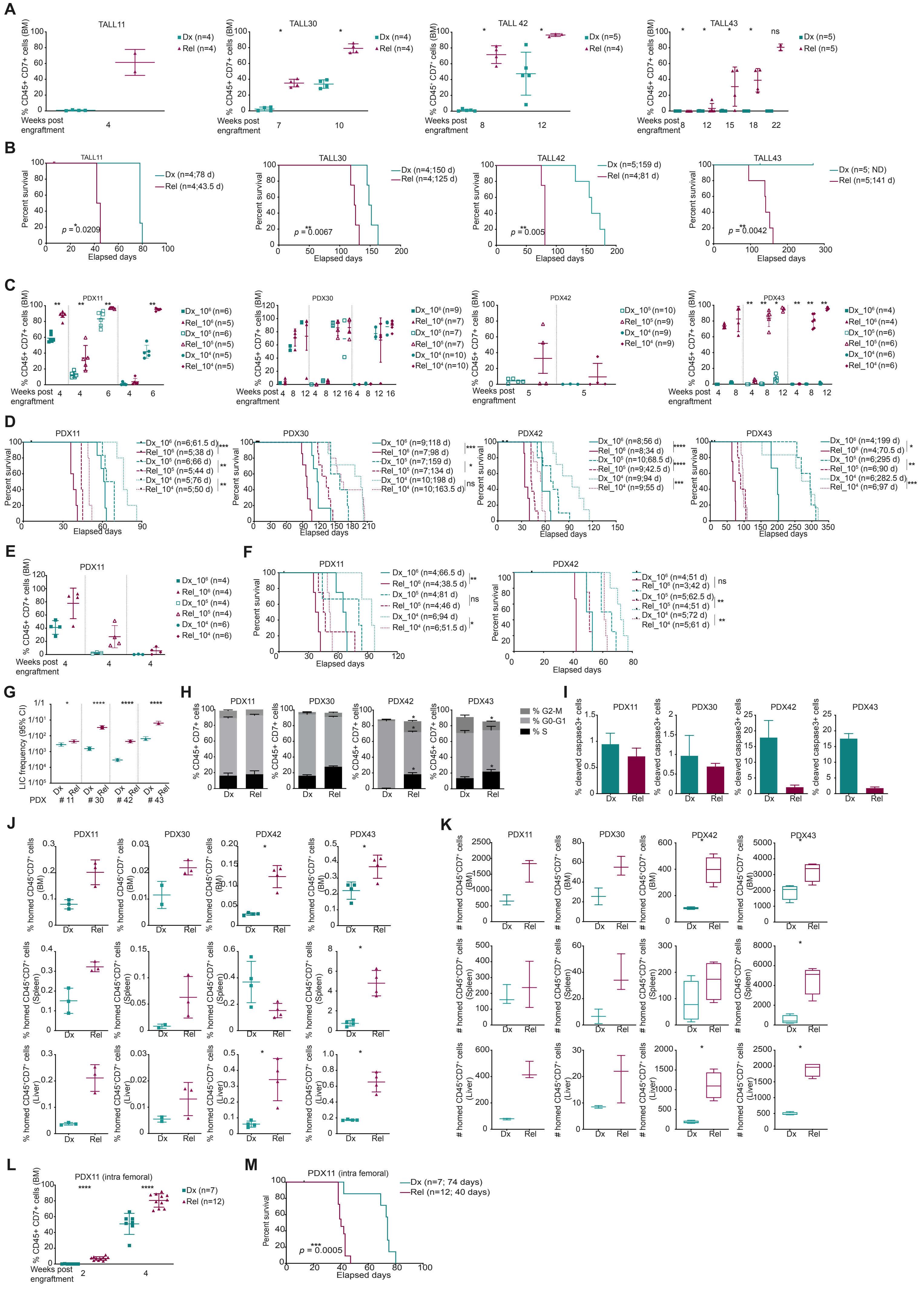

### Supplementary Figure 2

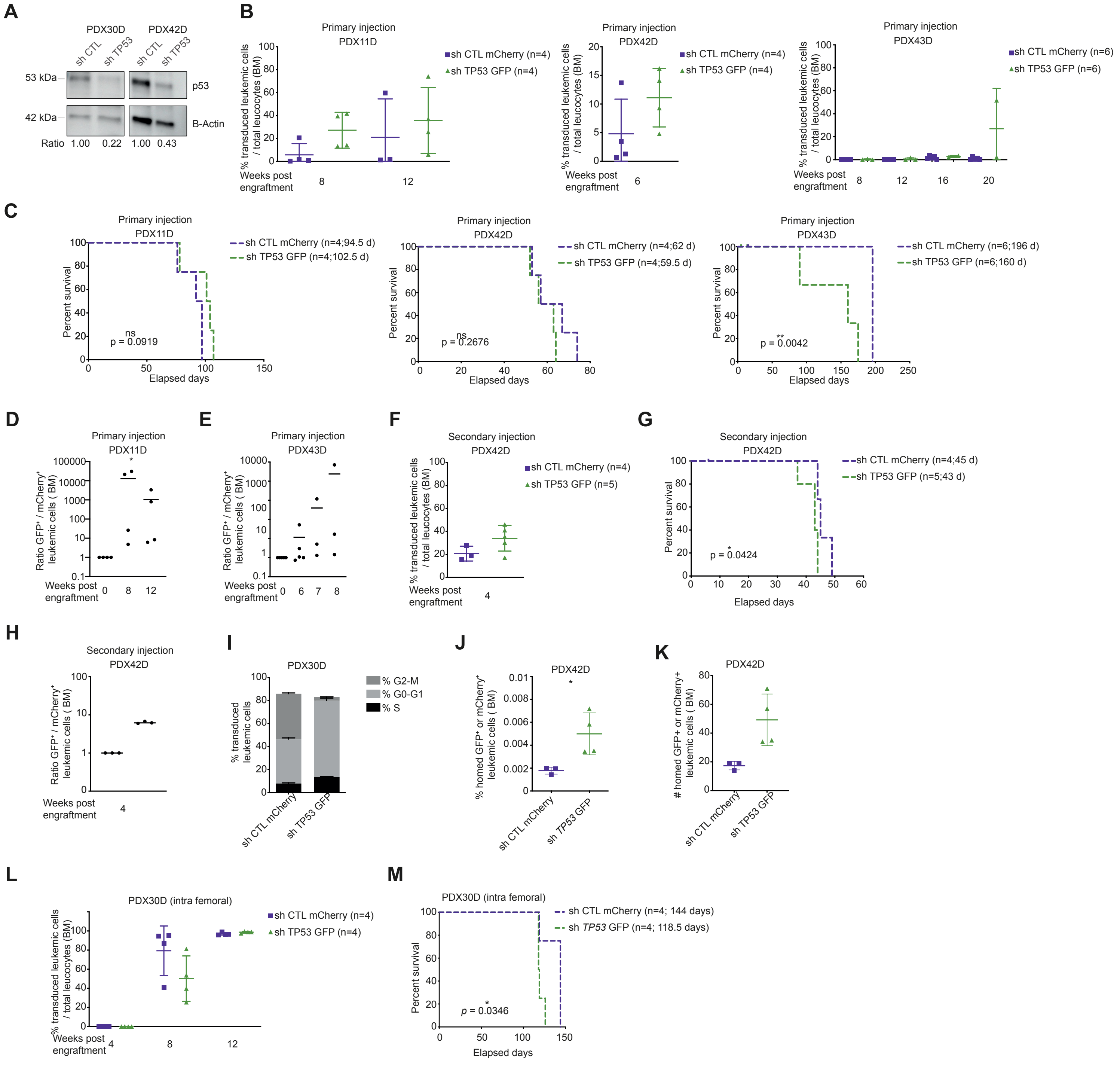

### Supplementary Figure 3

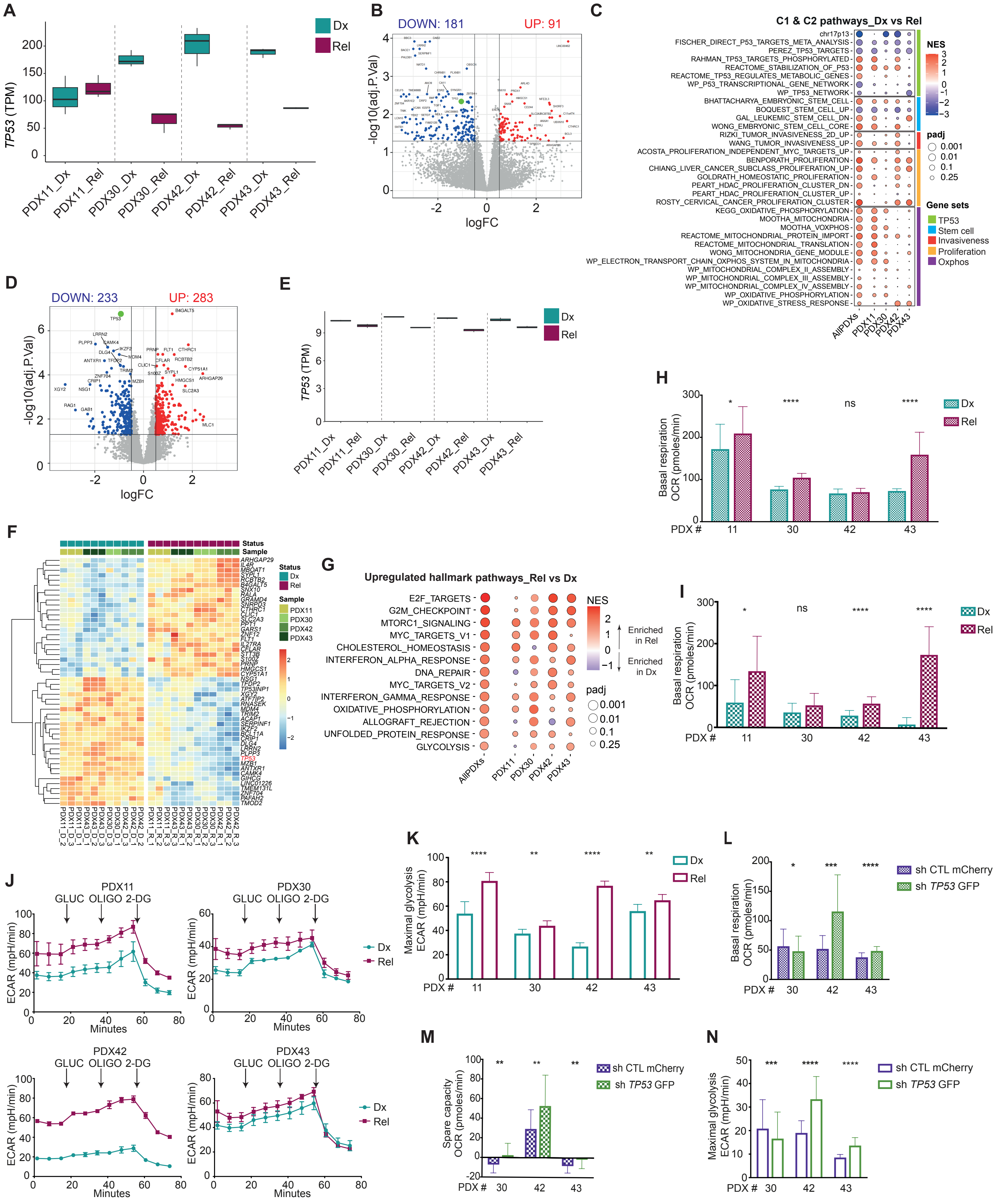

### Supplementary Figure 4

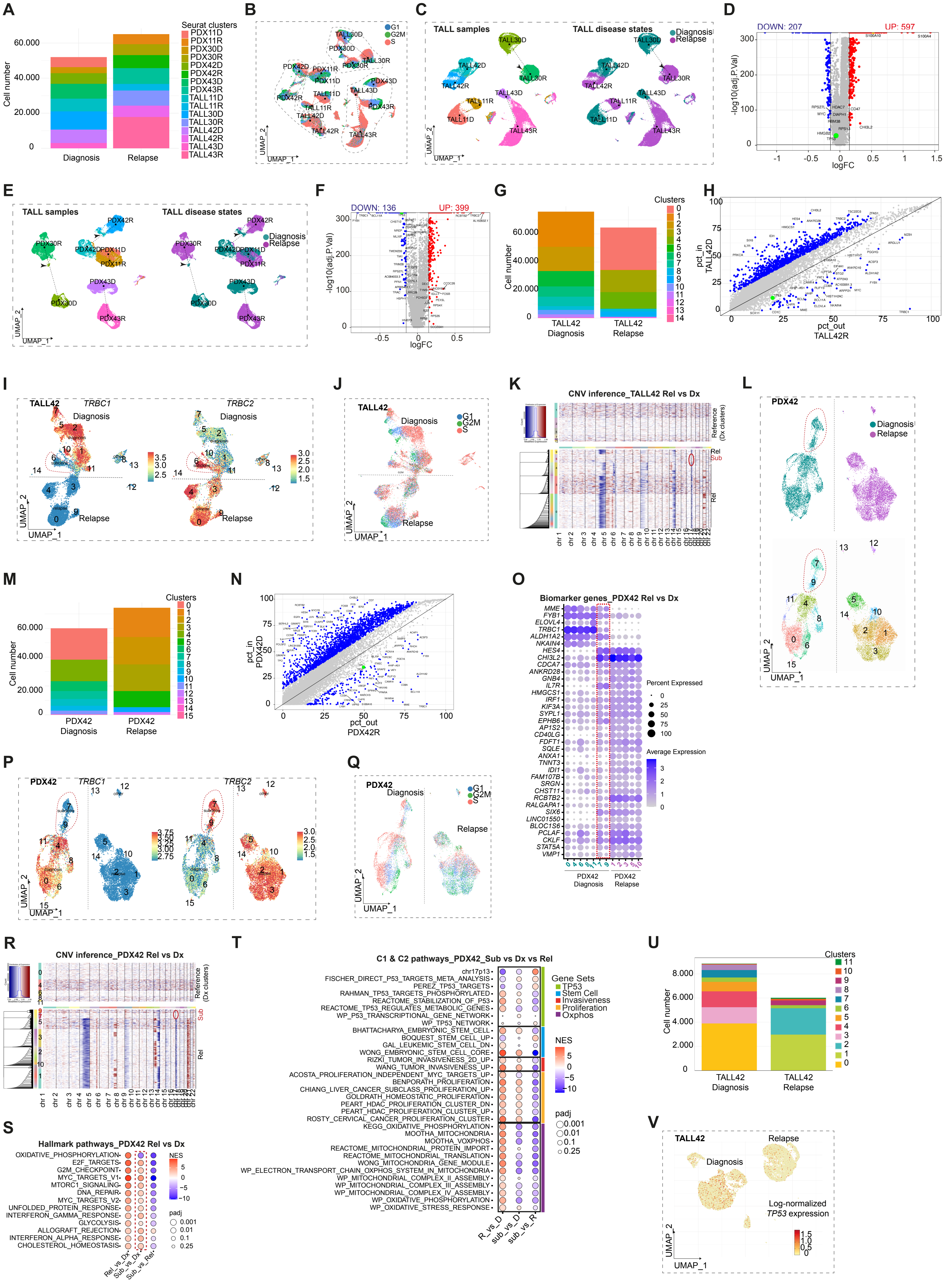

### Supplementary Figure 5

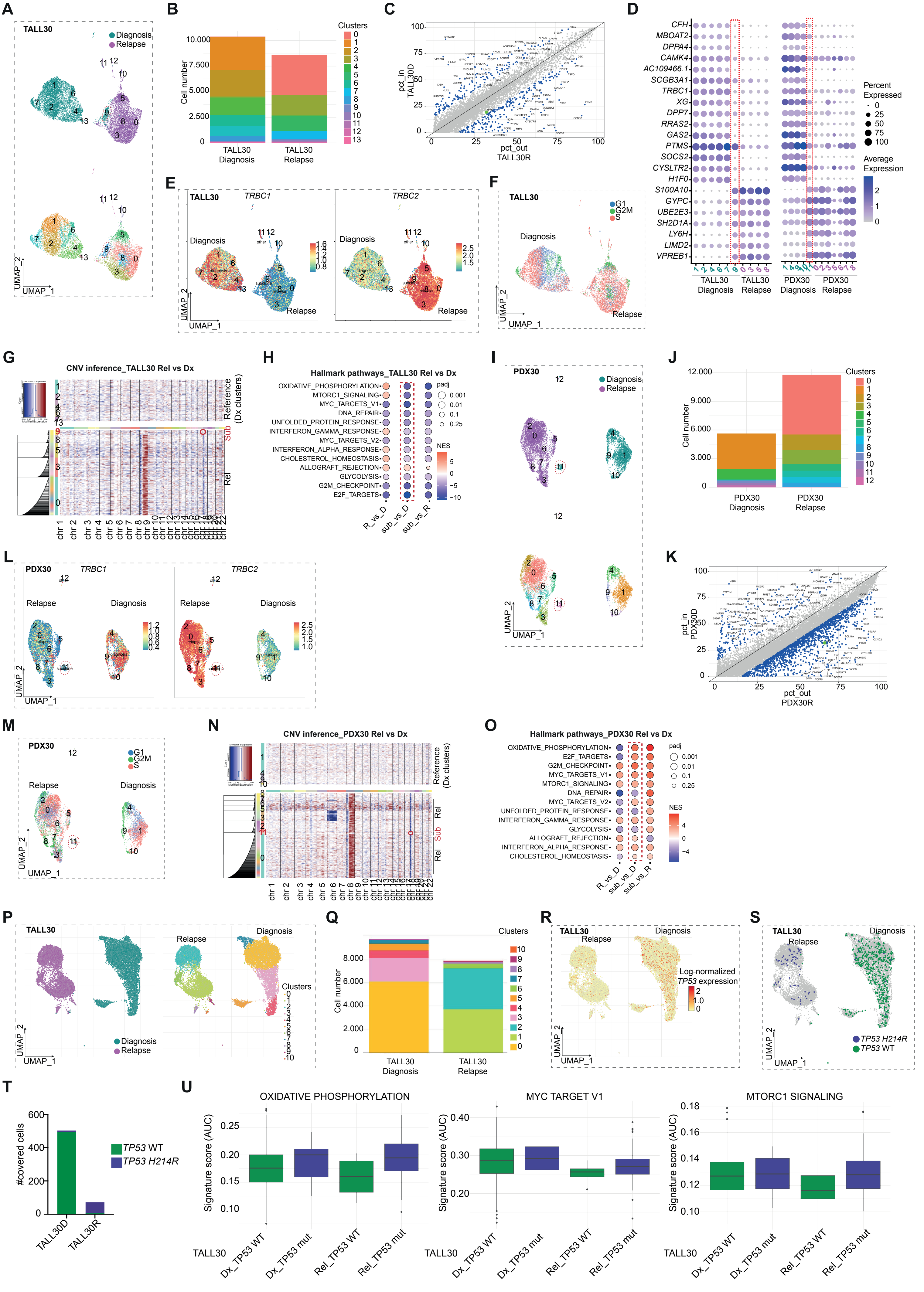
